# Multimodal anatomical mapping of subcortical regions in Marmoset monkeys using high-resolution MRI and matched histology with multiple stains

**DOI:** 10.1101/2023.03.30.534950

**Authors:** Kadharbatcha S Saleem, Alexandru V Avram, Cecil Chern-Chyi Yen, Kulam Najmudeen Magdoom, Vincent Schram, Peter J Basser

**Affiliations:** Section on Quantitative Imaging and Tissue Sciences (SQITS), Eunice Kennedy Shriver National Institute of Child Health and Human Development (NICHD), NIH, Bethesda, MD 20892; Center for Neuroscience and Regenerative Medicine (CNRM), Henry M. Jackson Foundation (HJF) for the Advancement of Military Medicine, Bethesda, MD 20817; National Institute of Neurological Disorders and Stroke (NINDS); Microscopy and Imaging Core (MIC)Eunice Kennedy Shriver National Institute of Child Health and Human Development (NICHD), NIH, Bethesda, MD 20892

**Author notes:** Correspondence to: Kadharbatcha Saleem, PhD, Section on Quantitative Imaging and Tissue Sciences (SQITS) Eunice Kennedy Shriver – National Institute of Child Health and Human Development (NICHD), National Institutes of Health (NIH), Bethesda, Maryland 20892.

**Keywords:** MAP-MRI, DTI, diffusion propagator, T2W, histology, subcortical, nuclei, fiber tracts, iron, macaque monkey, and marmoset monkey

## Abstract

Subcortical nuclei and other deep brain structures play essential roles in regulating the central and peripheral nervous systems. However, many of these nuclei and their subregions are challenging to identify and delineate in conventional MRI due to their small size, hidden location, and often subtle contrasts compared to neighboring regions. To address these limitations, we scanned the whole brain of the marmoset monkeys in *ex vivo* using a clinically feasible diffusion MRI method, called the mean apparent propagator (MAP)-MRI, along with T2W and MTR (T1-like contrast) images acquired at 7 Tesla. Additionally, we registered these multimodal MRI volumes to the high-resolution images of matched whole-brain histology sections with seven different stains obtained from the same brain specimens. At high spatial resolution, the microstructural parameters and fiber orientation distribution functions derived with MAP-MRI can distinguish the subregions of many subcortical and deep brain structures, including fiber tracts of different sizes and orientations. The good correlation with multiple but distinct histological stains from the same brain serves as a thorough validation of the structures identified with MAP-MRI and other MRI parameters. Moreover, the anatomical details of deep brain structures found in the volumes of MAP-MRI parameters are not visible in conventional T1W or T2W images. The high-resolution mapping using novel MRI contrasts, combined and correlated with histology, can elucidate structures that were previously invisible radiologically. Thus, this multimodal approach offers a roadmap toward identifying salient brain areas *in vivo* in future neuroradiological studies. It also provides a useful anatomical standard reference for the region definition of subcortical targets and the generation of a 3D digital template atlas for the marmoset brain research (Saleem et al., 2023). Additionally, we conducted a cross-species comparison between marmoset and macaque monkeys using results from our previous studies (Saleem et al., 2021). We found that the two species had distinct patterns of iron distribution in subregions of the basal ganglia, red nucleus, and deep cerebellar nuclei, confirmed with T2W MRI and histology.

## Introduction

The basal ganglia, thalamus, hypothalamus, brainstem nuclei, and amygdala are subcortical regions that regulate sensorimotor, cognitive, limbic, or autonomic (sympathetic and parasympathetic) functions. Continuing advances in neuroimaging have yielded a growing number of useful anatomical contrasts, in addition to the conventional T1- and T2-weighted MR images, thereby enabling multiparametric mapping of subcortical regions and white matter fiber tracts in humans and the non-human primate (NHP), such as the macaque monkey (Plantinga et al., 2014; Saleem et al., 2021). In particular, the diffusion MRI (dMRI) is a noninvasive preclinical and clinical neuroimaging method that provides volumetric images of deep brain/subcortical structures and enables visualization of associated white matter pathways with tractography methods (Calabrese et al., 2015; Oishi et al., 2020). dMRI is sensitive to the microscopic motions of water molecules diffusing in tissue and can therefore reflect features of cellularity or myelin content (Pierpaoli and Basser, 1996; Pierpaoli et al., 1996). Signal models of dMRI, such as diffusion tensor imaging (DTI), can measure diffusion in both isotropic and anisotropic tissue, such as white matter, and characterize the underlying microstructure with eloquent scalar parameters such as the fractional anisotropy (FA) and the mean diffusivity (MD) (Basser, 1995; Basser et al., 1994).

Mean apparent propagator (MAP)-MRI (Ozarslan et al., 2013) is a clinically feasible (Avram et al., 2016) advanced dMRI method that efficiently quantifies the 3D net displacements of water molecules diffusing in tissues, i.e., the diffusion propagators. In each voxel, the propagator describes comprehensively water diffusion processes in tissues with complex microstructures. MAP-MRI generalizes and subsumes other dMRI methods, such as DTI or diffusion kurtosis imaging (DKI) (Avram et al., 2017), and allows us to derive a wide range of dMRI microstructural parameters, including the mean, axial, and radial diffusivities (MD, AD, and RD, respectively), fractional anisotropy (FA), or the DEC maps (Pajevic and Pierpaoli, 1999). Moreover, MAP-MRI yields a family of new microstructural parameters that quantify important features of the diffusion propagators, such as the zero-displacement probabilities, non-gaussianity index (NG), and propagator anisotropy (PA) (Avram et al., 2014a; Avram et al., 2018a; Avram et al., 2017). Taken together, the scalar DTI/MAP parameters reflect features of the tissue microstructure more efficiently than conventional dMRI methods (Hutchinson et al., 2018), providing a richer contrast in both gray and white matter compared to conventional T1 or T2-weighted MRIs. Therefore, they are well-suited for detailed anatomical mapping of cortical and subcortical structures (Avram et al., 2022b; Saleem et al., 2021).

Several studies have delineated the subcortical structures in humans using high-resolution *in vivo* MRI with multiple image contrasts or *ex vivo* spin echo (SE) T2W MRI with histological stains (Abosch et al., 2010; Deistung et al., 2013a; Deistung et al., 2013b; Ewert et al., 2018; Hoch et al., 2019a; Hoch et al., 2019b; Keuken et al., 2014; Lenglet et al., 2012; Pauli et al., 2018; Rijkers et al., 2007). In contrast, only a few studies have performed the detailed mapping of subcortical regions using combined MRI and histology in the marmoset. Newman and colleagues have mapped the subcortical regions on a series of individual photographs of Nissl-stained sections with closely matched conventional T2W MR images obtained from a different marmoset brain (Newman et al., 2009). Similar histology-based mappings of subcortical regions have been published in book form with or without MRI (Hardman and Ashwell, 2012; Iriki et al., 2018; Palazzi and Bordier, 2008; Paxinos et al., 2012; Stephan et al., 1980; Yuasa et al., 2010). Concurrently, only a few studies have produced detailed mappings of subcortical regions in the macaque monkey using combined MRI and histology. For more details, see the introduction section in our previous publication (Saleem et al., 2021).

Because subcortical structures are usually concentrated in a small brain region and have limited contrast on MRI, imaging detailed neuroanatomy with conventional T1W and T2W scans alone without the corresponding matched histological information from the same animal is challenging and prone to errors. As shown in our previous study in the macaque monkey (Saleem et al., 2021), combining volumes of different MRI markers acquired with high-spatial resolution (200 μm), augmented by histological information derived from the same brain specimen, is pivotal in delineating nuclei and fiber tracts in deep brain structures, including substructures and laminae, e.g., in the basal ganglia and thalamus. Thus, a complete-3D brain segmentation based on MRI-histology correlations is critical for assisting neuroanatomical and neuroimaging studies in marmosets.

Here, we combined high-dimensional MRI data (DTI/MAP-MRI, T2W, and MTR) and the matched high-resolution images of histology sections with multiple stains derived from the same marmoset brain specimen to map the location, boundaries, and microarchitectural features of subcortical regions and associated white matter pathways in a complete marmoset brain *ex vivo*. This integrated multimodal approach yields a more objective and reproducible delineation of gray matter nuclei and their boundaries in the deep brain structures, which includes the basal ganglia, thalamus, hypothalamus, limbic region (amygdala), basal forebrain, and the rostrocaudal extent of the brainstem (midbrain, pons, and medulla).

## Materials and Methods

### Perfusion fixation

One adult male (case 1) and one female (case 2) marmoset monkey (*Callithrix jacchus*), weighing 340 gm and 283 gm, respectively, were perfused for the combined *ex vivo* MRI and histological studies. The animals were deeply anesthetized with sodium pentobarbital and perfused transcardially with 0.5 liters of heparinized saline, followed by 2 liters of 4% paraformaldehyde, both in 0.1 M phosphate buffer (pH 7.4). After perfusion, the brains were removed from the cranium, photographed, and post-fixed for 24 h in the same buffered paraformaldehyde solution and eventually transferred into 0.1 M phosphate-buffered saline (PBS) with sodium azide before MRI scanning. All procedures adhered to the Guide for the Care and Use of Laboratory Animals (National Research Council) and were carried out under a protocol approved by the Institutional Animal Care and Use Committee of the National Institute of Mental Health (NIMH) and the National Institutes of Health (NIH).

### *Ex vivo* MRI

#### Data acquisition

In preparation for the MRI experiments, we positioned the fixed brain specimen from case 1 in a custom 3D printed brain mold (Fig. 1) and then inside a custom 30 mm diameter cylindrical container. We filled the 3D mold and container with Fomblin and gently stirred them under a vacuum for several hours to remove potential air bubbles in and around the brain. Subsequently, we sealed the container and prepared the sample for MRI using a Bruker 7T/300 mm horizontal bore MRI scanner with a 30 mm inner diameter quadrature millipede coil (ExtendMR; http://www.extendmr.com/). We eventually registered all MRI volumes to a high-resolution multi-subject *in vivo* T1W anatomical template of the marmoset brain oriented to the stereotaxic coordinates (Liu et al., 2021) to generate the standard 3D digital template atlas (in progress). We adopted a similar procedure for the sample preparation before MRI scanning in cases 1 and 2 but used slightly different MRI scan parameters between these cases.

In case 1, we acquired MAP-MRI data with an isotropic resolution of 150 µm, i.e., a 288x184x184 imaging matrix on a 4.32×2.76×2.76 cm field-of-view (FOV), using a 3D diffusion spin-echo (SE) echo-planar imaging (EPI) sequence with 48 ms echo time (TE), 650 ms repetition time (TR), ten segments, and 1.33 partial Fourier acceleration. We obtained a total of 256 diffusion-weighted images (DWIs) using multiple b-value shells: 100, 500, 1000, 1500, 2500, 3500, 4500, 5250, 7000, 8500, 10000s/mm^2^ and diffusion-encoding gradient orientations (7, 10, 12, 15, 21, 22, 27, 31, 35, 36, and 40, respectively) uniformly sampling the unit sphere on each shell (Avram et al., 2018b; Koay et al., 2012). The diffusion gradient pulse durations and separations were δ=6 ms and Δ=28 ms, respectively. We also acquired a magnetization transfer (MT) prepared 3D gradient echo scan with 150 µm isotropic resolution (a 288x184x184 imaging matrix on a 4.32×2.76×2.76 cm FOV) with a 15° excitation flip angle and TE/TR=3.7/37ms. The parameters of the MT saturation pulse were 2 kHz offset, 12.5 ms Gaussian pulse with 6.74 µT peak amplitude, and 540° flip angle. We obtained four averages for each MT on and off scans. The total duration of the MAP-MRI scan was 85 h and 20 min, and the MT scan was 11 h and 8 min.

In case 2, we obtained MAP-MRI data with an isotropic resolution of 150 µm, i.e., a 256x160x160 imaging matrix on a 3.84×2.40×2.40 cm FOV, using a 3D diffusion spin-echo (SE) echo-planar imaging (EPI) sequence with 43 ms TE, 1400 ms TR, eight segments and 1.25 partial Fourier acceleration. We acquired a total of 112 DWIs using multiple b-value shells: 100, 1000, 2500, 4500, 7000, and 10000 s/mm^2^ and diffusion-encoding gradient orientations (3, 9, 15, 21, 28, and 36, respectively) uniformly sampling the unit sphere on each shell (see related references in the previous paragraph). Two averages were acquired for the scan with b=10000 s/mm^2^ to improve SNR. The diffusion gradient pulse durations and separations were δ=8 ms and Δ=20 ms, respectively. We also acquired a MT-prepared 3D gradient echo scan with 150 µm isotropic resolution (a 256x160x160 imaging matrix on a 3.84×2.40×2.40 cm FOV), a 15° excitation flip angle, TE/TR=3.7/37ms. The parameters of the MT saturation pulse were 2 kHz offset, 12.5 ms Gaussian pulse with 6.74 µT peak amplitude, and 540° flip angle. We obtained four averages for each MT on and off scans. The total duration of the MAP-MRI scan was 74 h and 40 min, and the MT scan was 8 h and 31 min.

#### Data processing

The post-processing and analysis of data are similar in both cases. We processed all DWIs using the TORTOISE software packages (Pierpaoli et al., 2010) and the MT ratio (MTR) image as the structural template to correct for Gibbs ringing artifacts, intrascan motion (drift), and imaging distortions due to magnetic field inhomogeneities and diffusion gradient eddy currents. We estimated the mean apparent propagator in each voxel by fitting the diffusion data with a MAP series approximation, truncated at order 4, using a MATLAB implementation. We computed DTI parameters, such as the mean diffusivity (MD), fractional anisotropy (FA), axial diffusivity (AD), and radial diffusivity (RD), as well as MAP-MRI tissue parameters, such as the return-to-origin probability (RTOP), return-to-axis probability (RTAP), return-to-plane probability (RTPP), propagator anisotropy (PA), non-gaussianity (NG) and the non-diffusion attenuated (amplitude) image, which provides a T2W contrast. We also estimated in each voxel the orientation distribution function (ODF) and fiber ODFs (fODFs) and visualized them with the MrTrix3 software package (Tournier et al., 2012). From the DTI model, we computed the linear, planar, and spherical anisotropy coefficients, CL, CP, and CS, respectively, that characterize the shape of the underlying diffusion tensor (Westin et al., 2002). The MT ratio (MTR) was computed from the images acquired with and without MT preparation. The high contrast between gray and white matter (GM and WM, respectively) in the MTR image allowed reliable image registration to the DWIs.

We also acquired a high-resolution T2W scan from case 2 without the brain mold, using a 7T vertical wide bore Bruker MRI scanner with a microimaging probe having a 25 mm inner diameter quadrature coil. The dataset is a 100-μm isotropic 3D T2 weighted fast spin echo sequence with a TR = 500 ms, effective TE = 60 ms, RARE factor = 8, readout BW = 50 kHz, 36 mm x 24 mm x 24 mm FOV, and eight averages. The total duration of this scan was 8 h. We used this 3D dataset to delineate deep brain nuclei in the brainstem and confirmed the structures with different histological stains, including Perls Prussian blue (iron) stain (see Figs. 10, 13, 14). In addition to our multiple datasets from cases 1 and 2, we also analyzed previously published and publicly shared marmosets high-resolution diffusion MRI volumes (Liu et al., 2020) to delineate the brainstem nuclei and fiber tracts in a sagittal plane of sections (see supplement Fig. 1).

##### Histological processing specific to case 1

Following MRI acquisition, the intact brain specimen from case 1 was prepared for histological processing with five different stains. All histological processing (section cutting and staining) was done by FD NeuroTechnologies, Columbia, Maryland. The formaldehyde-perfusion-fixed marmoset brain was cryoprotected with FD tissue cryoprotection solution™ (FD NeuroTechnologies, Columbia, MD) for 15 days and then rapidly frozen in isopentane pre-cooled to -70°C with dry ice. The frozen brain was stored in a -80°C freezer before sectioning. Note that we did not cut the brain into blocks before freezing. Serial frozen coronal sections (50-µm thick) were cut on a sliding microtome through the entire brain as one tissue block, including the cerebrum, brainstem, and cerebellum. We collected a total of 670 sections from this brain and sorted them into five interleaved parallel series (134 sections per set), and processed them for different cell bodies and fiber stains (Fig. 1). All the sections were stained in this animal (see below). We also collected block-face images of the frozen tissue block at every 250 μm interval.

**Fig. 1.**
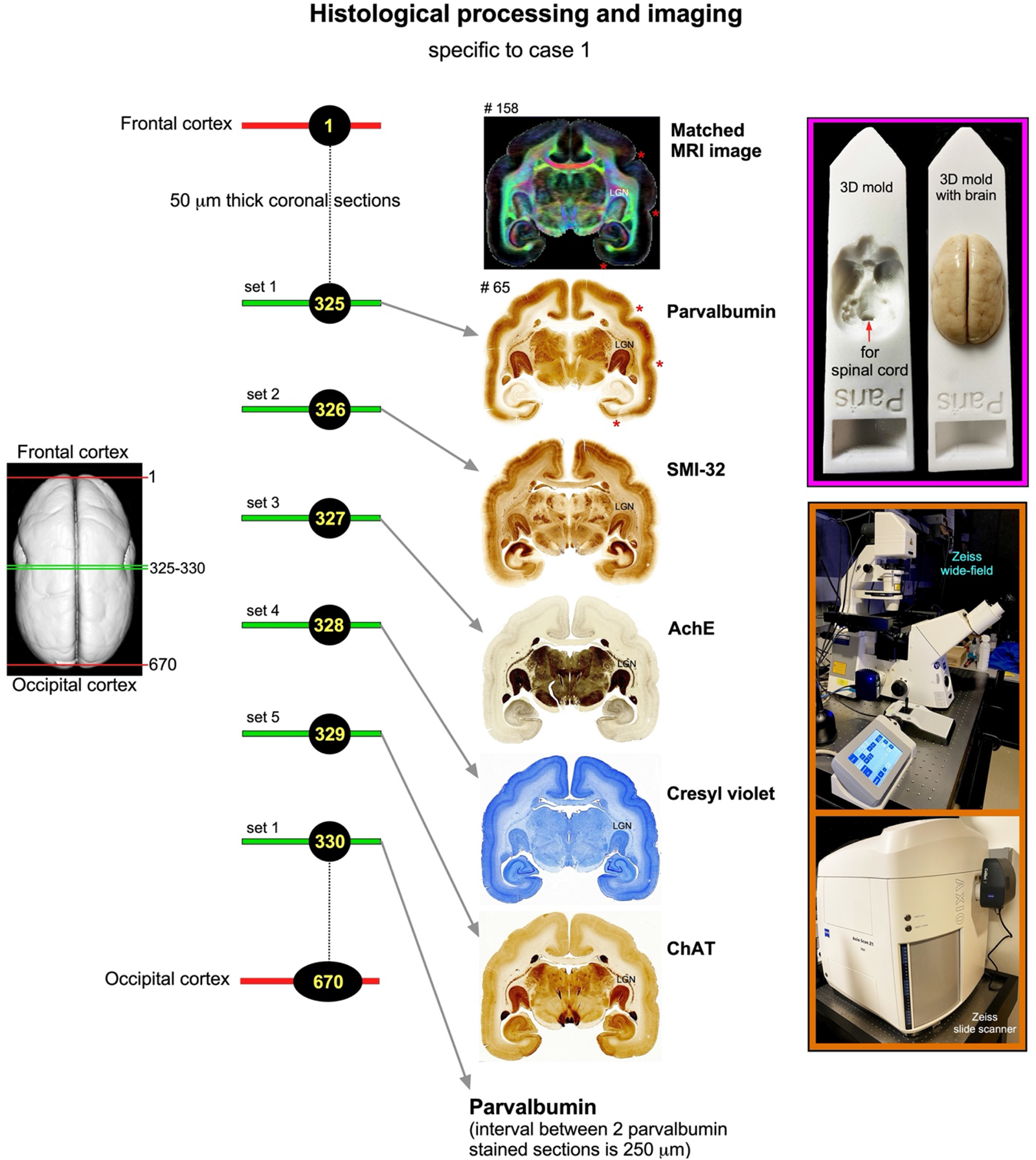
Histological staining and high-resolution imaging in case 1. Frozen sections were cut coronally from the frontal cortex to the occipital cortex at 50μm thickness on a sliding microtome. In total, 670 sections were collected, and all the sections were processed with different cell bodies and fiber stains. This example shows an adjacent series of five sections at the level of the anterior temporal cortex (325-329) stained with parvalbumin, SMI-32, AchE, Cresyl violet, and ChAT, respectively. The rostrocaudal locations of these sections are shown on the rendered brain image on the left. This sequence of staining is followed from the frontal to the occipital cortex. We obtained 134 stained sections in each series, and the interval between two adjacent sections in each series is 250 μm. The high-resolution images of stained sections were captured using a Zeiss wide-field microscope and a Zeiss Axioscan Z1 high-resolution slide scanner (inset on the bottom right). These histology images were then aligned manually with the corresponding MAP-MRI (top) and other MRI parameters of the same specimen to allow visualization and delineation of subcortical structures in a specific region of interest. In both MRI and histology sections, note the correspondence of sulci (red stars), gyri, and deep brain structures (e.g., LGN-lateral geniculate nucleus). #65 refers to the section number in each set/series of stained sections, and #158 indicates the matched MRI slice number in 3D volume. The unique characteristics of each stain are described in materials and methods. The inset on the top right shows the 3D brain mold with or without the brain specimen for MR imaging. For other details, see the “data acquisition” section above.

**Fig. 2.**
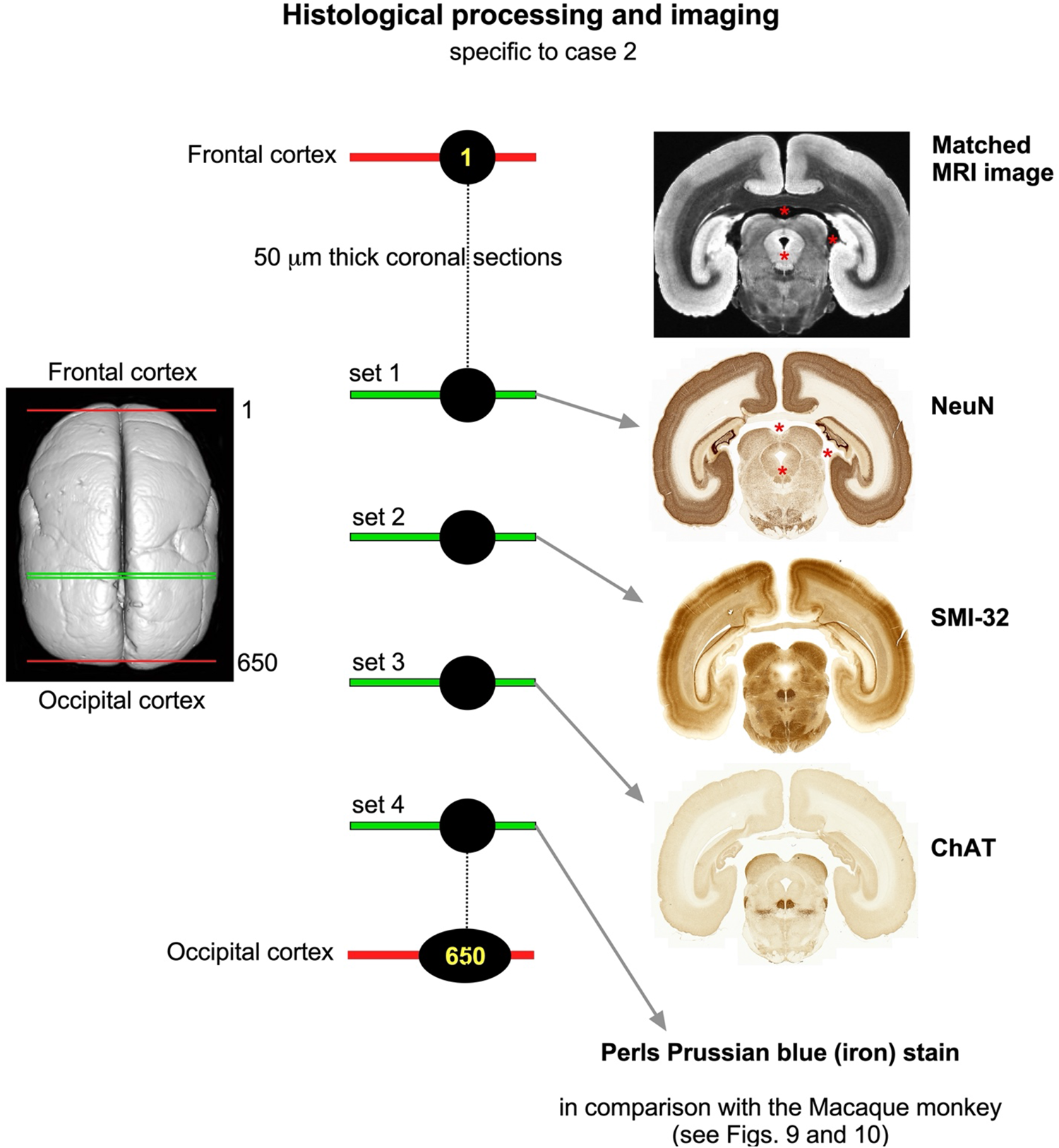
Histological staining and high-resolution imaging in case 2. Like case 1, the frozen sections were cut coronally from the frontal cortex to the occipital cortex at 50μm thickness on a sliding microtome. In total, 650 sections were collected and divided into ten series. The first three sets, with 65 sections each, were stained immunohistochemically with NeuN, SMI-32, and ChAT, respectively. The selected sections from the fourth set were stained histochemically with Perls Prussian blue. The high-resolution images of stained sections were captured using a Zeiss Axioscan Z1 high-resolution slide scanner (see Figure 1). These histology images were then aligned manually with the corresponding MRI of the same brain specimen to allow visualization and delineation of subcortical structures in a specific region of interest. Note the correspondence of anatomical landmarks in both MRI and histology sections (red stars). For other details, see figure 1 legend.

#### Immunohistochemistry

The sections of 1^st^, 2^nd^, and 5^th^ sets were processed for parvalbumin (PV)-, neurofilament (SMI 32)-, and choline acetyltransferase (ChAT)-immunohistochemistry with the commercially available antibodies (see below; Fig. 1). Briefly, after inactivating endogenous peroxidase activity with hydrogen peroxidase, sections were incubated free-floating in 0.01 M phosphate-buffered saline (PBS, pH 7.4) containing 0.3% Triton X-100 (MilliporeSigma, St. Louis, MO), 1% normal blocking serum (Jackson ImmunoResearch, West Grove, PA), and one of the following primary antibodies: mouse monoclonal anti-Parvalbumin antibody (Cat. # P3088, 1:3,000, MilliporeSigma), mouse monoclonal anti-nonphosphorylated neurofilament H (clone SMI 32, Cat. # 801701, 1:1,000, BioLegend, San Diego, CA), and goat anti-choline acetyltransferase antibody (Cat. # AB144P, 1:400, MilliporeSigma) for 67 h at 4°C. The immunoreaction products were then visualized according to the avidin-biotin complex method (Hsu et al., 1981) using the Vectastain elite ABC kit (Vector Lab., Burlingame, CA) and 3’,3’-diaminobenzidine (MilliporeSigma, St. Louis, MO) as a chromogen. After thorough washes, all sections were mounted on gelatin-coated microscope slides, dehydrated in ethanol, cleared in xylene, and coverslipped with Permount® (Fisher Scientific, Fair Lawn, NJ, USA).

#### Histochemistry

The sections of the 4^th^ set were mounted on gelatin-coated microscope slides and stained with the FD cresyl violet solution™ (FD NeuroTechnologies) for Nissl substance. The sections of the 3^rd^ set were processed with the modified acetylcholinesterase (AchE) histochemistry method (Naik, 1963). Briefly, after washing in PBS, sections were pretreated with 0.05 M acetic buffer (pH 5.3) 3 times, 3 min each. Sections were incubated free-floating in solution A containing sodium acetate, copper sulfate, glycine, ethopropazine, and acetylthiocholine for three days. Sections were then incubated in solution B containing sodium sulfide, followed by incubation in solution C containing silver nitrate. Subsequently, sections were re-fixed in 0.1 M phosphate buffer containing 4% paraformaldehyde overnight. All the above steps were carried out at room temperature, and each step was followed by washes in distilled water. After thorough washes in distilled water, sections were mounted on gelatin-coated slides. Following dehydration in ethanol, the sections were cleared in xylene and coverslipped with Permount® (Fisher Scientific).

##### Histological processing specific to case 2

The preparation of this brain specimen for the histological work is the same as in case 1. In addition to SMI-32 and ChAT, we stained the brain sections from case 2 with new markers: the NeuN (neuron-specific nuclear protein) and the Perls Prussian blue or iron stain (Fig. 2). The immunohistochemical staining methods used for SMI-32 and ChAT are comparable to case 1 (see above).

#### NeuN Immunohistochemistry

Briefly, after inactivating endogenous peroxidase activity with hydrogen peroxidase, sections were incubated free-floating in 0.01 M phosphate-buffered saline (PBS, pH 7.4) containing 0.3% Triton X-100 (MilliporeSigma, St. Louis, MO), 1% normal blocking serum (Jackson ImmunoResearch, West Grove, PA), and the following primary antibody for NeuN: (Cat. MAB377, 1:10,000, MilliporeSigma) for 67 h at 4°C. The immunoreaction products were then visualized according to the avidin-biotin complex method (Hsu et al., 1981) using the Vectastain elite ABC kit (Vector Lab., Burlingame, CA) and 3’,3’-diaminobenzidine (MilliporeSigma, St. Louis, MO) as a chromogen. After thorough washes, all sections were mounted on gelatin-coated microscope slides, dehydrated in ethanol, cleared in xylene, and coverslipped with Permount® (Fisher Scientific, Fair Lawn, NJ, USA). For Perls Prussian blue (iron) staining procedures, see the “Marmoset versus macaque” section below.

The histological stains used in this study labeled different types of neuronal cells- or both cell bodies and fiber bundles in cortical and subcortical regions. The calcium-binding protein parvalbumin (PV) was thought to play an essential role in intracellular calcium homeostasis. The antibody against PV has been shown previously to recognize a subpopulation of non-pyramidal neurons (GABAergic) in the neocortex and different types of neurons in subcortical structures (Jones, 1998; Jones and Hendry, 1989; Saleem et al., 2007). The SMI-32 antibody recognizes a non-phosphorylated epitope of neurofilament H (Goldstein et al., 1987; Sternberger and Sternberger, 1983) and stains a subpopulation of pyramidal neurons and their dendritic processes in the monkey neocortex (Hof and Morrison, 1995; Saleem and Logothetis, 2012). It is also a valuable stain for a vulnerable subset of pyramidal neurons in the visual, temporal, and frontal cortical areas visualized in postmortem brain of Alzheimer’s disease cases (Hof et al., 1990; Hof and Morrison, 1990; Thangavel et al., 2009). The SMI-32 can also detect axonal pathology in TBI brains (Johnson et al., 2016). The antibody against ChAT recognizes cholinergic neurons and has been a valuable stain for motor neurons in the monkey and human brainstem (e.g., cranial nerve nuclei, (Horn et al., 2018). AchE is an enzyme that catalyzes the breakdown of acetylcholine and is a valuable marker for delineating different cortical areas (Carmichael and Price, 1994) and major subcortical nuclei in the thalamus and brainstem (Horn et al., 2018; Jones, 1998). The NeuN (neuron-specific nuclear protein) antibody is the marker for various types of neuronal cell bodies in the cortical and subcortical areas (Mullen et al., 1992) and can be used to study neuronal differentiation and development.

#### Data analysis

The high-resolution images of all stained sections were captured using a Zeiss wide-field microscope and Zeiss high-resolution Axioscan-Z1 slide scanner at 5X objective, and these digital images were adjusted for brightness and contrast using Adobe Photoshop (24.2 v). These images were then aligned manually with the corresponding images of DTI/MAP parameters along with the estimated T_2_-weighted (i.e., non-diffusion weighted) and the MTR images to allow visualization and delineation of subcortical structures in specific region-of-interest (ROIs) (e.g., Figs. 4, 5, 11). Some structures, like striatum and pallidum, were demarcated by comparing them side-by-side on the matched MRI and histology sections, but for others (e.g., amygdala), we used a different approach to outline their subregions as described in our previous study (Saleem et al., 2021). In brief, we first superimposed a histology section onto the matched MRI slice and then manually rotated and proportionally scaled it to match the boundaries of the different nuclei and their subregion on the MRI using the transparency function in Canvas X Draw software. Finally, the borders of the subregions were manually traced on the histology sections and translated onto the superimposed underlying MR images using the polygon-drawing tools with smooth and grouping functions in this software (e.g., Figs. 10, 15).

The histological sections were matched well with the MRIs in this study. It was not necessary to resample the MRI volume to achieve good alignment with the histological images in the current study. The alignment of these images was only possible by carefully orienting the entire brain specimen with reference to sagittal MRI images from this specimen. These steps were done on the microtome stage before attaching and freezing the brain specimen with dry ice. These steps enabled us to match the sulci, gyri, and region of interest (ROI) in deep brain structures in both MRI and histological sections, as shown in Figure 1 and the result section below (e.g., Fig. 11).

### Marmoset versus macaque

We found distinct MR Signal intensity differences in some subcortical regions between the macaque and the marmoset monkeys. For example, in T2W images, the basal ganglia and the deep cerebellar nuclei subregions exhibited significantly more hypointense signals in the macaque than in the marmoset. MRI studies in humans and nonhuman primates have interpreted and confirmed this hypointense signal to a high level of intracellular ferric iron deposit (see results). To confirm this finding histologically, we used unstained sections matched with MRI slices from the same macaque and marmoset (case 2) brains for the iron or Perls Prussian blue stain. Since we had processed all the histology sections in marmoset case 1 for five different stains (see Fig. 1), we did not have an additional set of unstained sections from this case for iron staining. However, we used another perfusion-fixed marmoset brain (case 2) for MRI and histology work with multiple stains, including Perls Prussian blue stain for the ferric iron (see Fig. 10).

### Prussian blue histochemistry in the macaque and marmoset monkeys

Selected 50-µm thick free-floating sections cut previously from a rhesus macaque (Saleem et al., 2021) and the current marmoset monkey (case 2) brains were processed for Prussian blue histochemistry. In brief, after mounting sections on 50x75mm gelatin-coated microscope slides (1 section per slide), the sections were incubated in a freshly prepared aqueous solution containing 1% hydrochloric acid (Sigma, St. Louis, MO) and 1% potassium ferrocyanide (Sigma, St. Louis, MO) for 30 min at room temperature. After thorough washes in distilled water, sections were dehydrated in ethanol, cleared in xylene, and coverslipped in Permount^®^ (Fisher Scientific, Fair Lawn, NJ).

## Results

Using a combination of MRI and histology, we mapped deep brain nuclei and their subregions, including the basal ganglia, thalamus, hypothalamus, brainstem (midbrain, pons, and medulla), amygdala, bed nucleus of stria terminalis, and the basal forebrain. We also included the architectonically and functionally distinct non-subcortical regions, such as different lobules of the cerebellar cortex and the hippocampal formation. In addition, we distinguished fiber tracts of different sizes and orientations associated with the basal ganglia, thalamus, brainstem, and cerebellum. Although we delineated and segmented 249 subcortical regions to create a 3D digital template atlas (in progress), it is beyond the scope of this study to describe the detailed anatomy and illustrate all the identified areas in this report. In the following sections, we first describe the spatial location, boundaries, and microarchitectural features of selected subcortical regions and associated white matter fiber pathways, identified with MRI markers and confirmed these structures in multiple histological stains, each of which has provided valuable distinctions between subregions. In addition, we also used the macaque MRI and histology data from our previous study (Saleem et al., 2021) to compare some deep brain structures with the marmoset brain. The data described in the results section is based on the marmoset cases 1 and 2, and the images derived from MRI and histology are comparable in both cases. The case numbers are indicated directly on the figures or in figure legends.

### MRI markers (Fig. 3)

MAP-MRI and other MRI parameters showed different gray and white matter contrast outside the cerebral cortex. In particular, PA or PA with fiber orientation distribution functions (fODFs) and the corresponding direction encoded color (DEC) map (Pajevic and Pierpaoli, 1999), RD, NG, RTAP, as well as T2W images revealed sharp boundaries and high contrast in the deep brain structures, resulting in a clear demarcation of anatomical structures such as nuclei/gray matter regions (e.g., medial dorsal nucleus of thalamus-MD, lateral geniculate nucleus-LGN, substantia nigra-SN, prerubral field-prf) (Fig. 3). All qualitative findings observed in MRI were confirmed using adjacent and matched histology sections with several stains (see materials and methods). For convenience, we combined both terminologies: fODFs derived directionally encoded color (DEC) map and abbreviated as DEC-FOD in the text and figures.

**Fig. 3.**
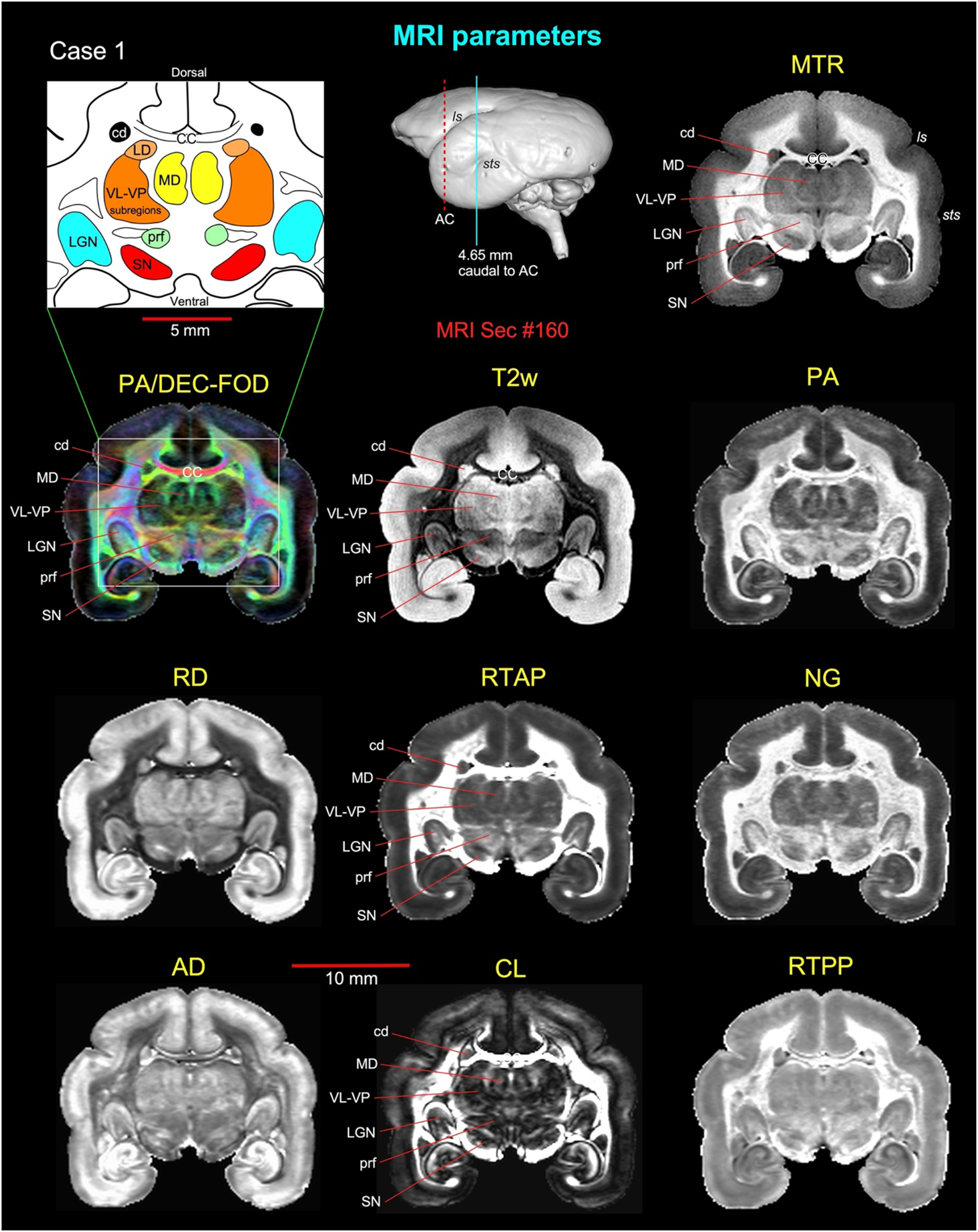
Subcortical regions in different MRI parameters. Matched coronal MR images from eight MAP-MRI parameters, T2-weighted (T2W), and the magnetization transfer ratio (MTR) images show selected subcortical regions: thalamic subregions (MD, LD, VL, VP, LGN), basal ganglia subregions (SN, cd), and a prerubral region (prf) anterior to the red nucleus. These areas are also illustrated in the corresponding drawing from the PA/DEC-FOD image on the top left (white box/inset). For more details on the thalamic subregions, see the result section. This MRI slice is located at the level of the rostral temporal cortex and 4.65 mm caudal to the anterior commissure (AC), as illustrated by a blue vertical line on the lateral view of the 3D rendered brain image from this case. Note that the contrast between these subcortical areas is distinct in different MRI parameters. ***Abbreviations: MAP-MRI parameters*:** AD-axial diffusivity; CL-linearity of diffusion from the tortoise; DEC-FOD-directionally encoded color-fiber orientation distribution; NG-non-gaussianity; PA-propagator anisotropy; RD-radial diffusivity; RTAP-return to axis probability; RTPP-return to plane probability. ***Subcortical regions:*** CC-corpus callosum; cd-caudate nucleus; LD-lateral dorsal nucleus; LGN-lateral geniculate nucleus; MD-medial dorsal thalamic nuclei; prf-prerubral field; SN-substantia nigra; subregions of VL-ventral lateral and VP-ventral posterior nuclei. ***Sulci:*** ls-lateral sulcus; sts-superior temporal sulcus.

### Delineation of selected subcortical gray and white matter regions

#### Basal ganglia

##### Striatum (**Fig. 4**)

The basal ganglia are composed of four major groups of nuclei: the striatum (caudate nucleus-cd, putamen-pu, and ventral striatum), pallidum, subthalamic nucleus (STN), and substantia nigra (SN). The spatial location, overall extent, and neighboring relationship between these structures can be appreciated in different views in 3D, as shown in Figure 4E, F. Both cd and pu exhibited similar hypo-or hyperintense contrast in MAP-MRI (PA/DEC-FOD), T2W, and MTR images, but this contrast is strikingly the opposite to the signal intensity found in the surrounding white matter regions, such as the anterior limb of the internal capsule (ica), corpus callosum (CC), and external capsule (ec) (Fig. 4A; T2W and MTR images are not shown in this figure). These structures’ location and well-defined contour, including bridges connecting cd and pu in MRI, are matched well with the histology sections stained with the ChAT (Fig. 4B), SMI-32, and PV. The ventral striatum (Haber, 2003; Heimer et al., 1982) includes the nucleus accumbens (NA) and is distinguished into the core (NAc) with hyperintense contrast and the shell (NAsh) with hypointense contrast in PA/DEC-FOD images (Fig. 4A). These NAc and NAsh subregions with light and dark contrast, respectively coincided with the light and darkly stained neuropil in these subregions in ChAT stained section (Fig. 4B). In addition, all subregions of the striatum reveal mosaic-like patterns with bright and dark contrasts in several MRI parameters but are most prominent in PA/DEC-FOD images, and might correspond to the light (“P”-patch or striosomes) and moderately-stained neuropil (“M”-matrix) regions, respectively in ChAT (Fig. 4C, D). The size and orientation of bright contrast regions in MRI did not match closely with lightly stained Patch compartments observed in adjacent ChAT stained sections.

**Fig. 4.**
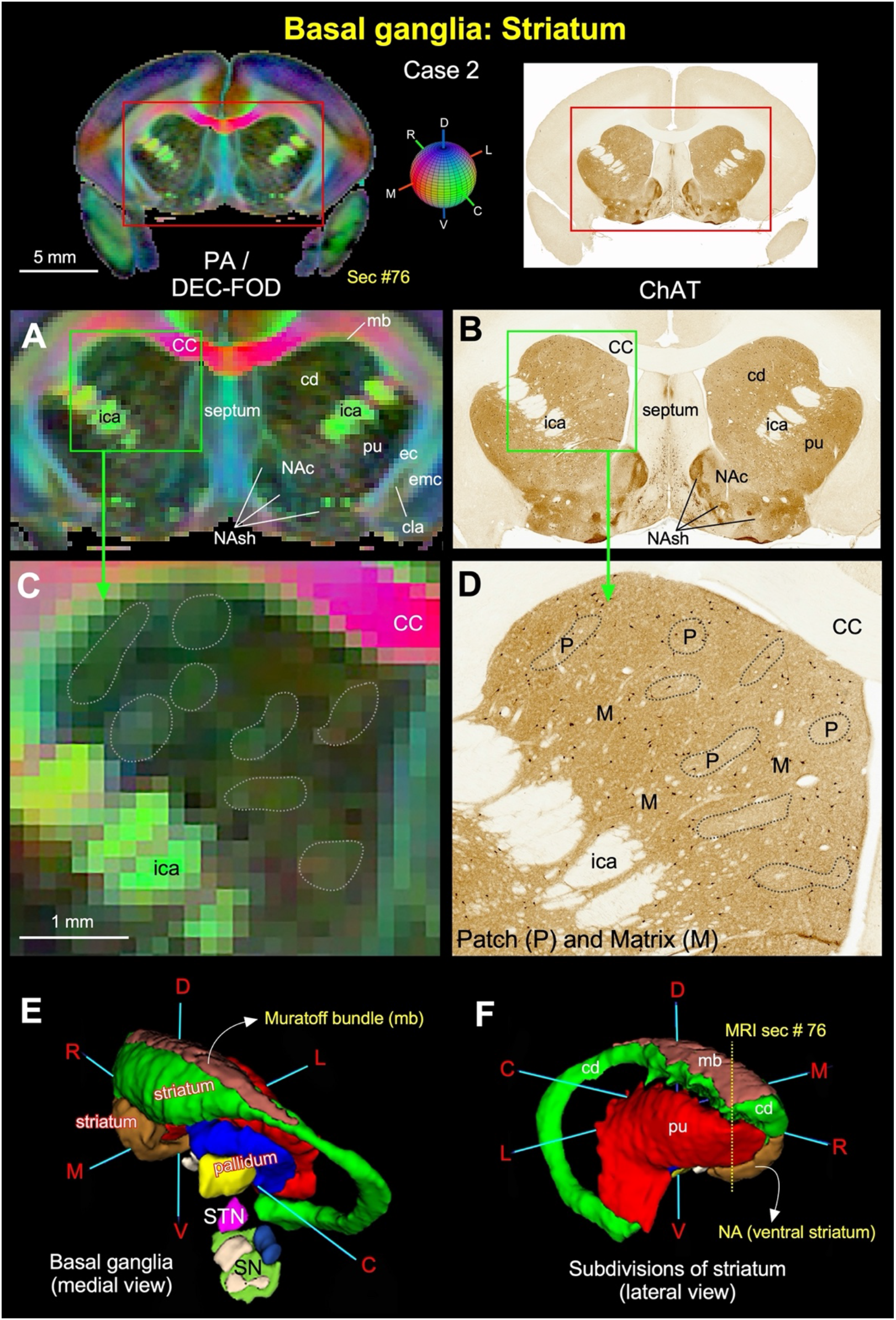
Striatum. **(A, B)** The zoomed-in view from the coronal sections on the top panel (red rectangles) shows the subregions of the striatum (caudate and putamen) and ventral striatum (nucleus accumbens and its subregions, core-NAc and shell-NAsh) in MAP-MRI (PA/DEC-FOD) and corresponding matched histology section stained for ChAT in case 2. Note the sharp contrast of these subregions from the surrounding white matter structures in MRI that corresponded well with the architectonic regions in histology sections, including the bridges connecting the caudate and putamen (compare A and B). **(C, D)** The high-power images of the head of the caudate nucleus (cd; green box) from DEC-FOD and adjacent ChAT stained sections show mosaic-like patterns with light and dark compartments (dashed outlined regions). These regions correspond to neurochemically defined compartments called patch (P) or striosomes and matrix (M) in ChAT stained section (see discussion). The “P” regions in ChAT (D) did not match with size and shape of the bright regions on the corresponding PA/DEC-FOD images. **(E, F)** Illustrate the spatial location and overall extent of the striatum with other basal ganglia subregions in 3D, reconstructed through serial MRI sections using ITK-SNAP. #76 in F indicates the corresponding rostrocaudal location of the coronal MRI slice illustrated on the top. ***Abbreviations:*** CC-corpus callosum; cd-caudate nucleus; cla-claustrum; ec-external capsule; emc-extreme capsule; ica-internal capsule, anterior limb; mb-Muratoff bundle; NAc-nucleus accumbens, core region; NAsh-nucleus accumbens, shell region; P-patch or striosomes; pu-putamen; SN-substantia nigra; STN-subthalamic nucleus. ***Orientation on 3D:*** D-dorsal; V-ventral; R-rostral; C-caudal; M-medial; L-lateral.

##### Pallidum (Fig. 5)

The pallidum is divided into the external and internal segments of the globus pallidus (GPe and GPi, respectively) and the ventral pallidum (VP) (Parent, 1990) that are readily identifiable as hyperintense in MTR or hypointense contrast in T2W images (Fig. 5). These pallidal subregions are easily delineated from the laterally adjacent putamen (pu), which shows the opposite signal intensity in both MR images (i.e., light versus dark contrast or vice-versa; Fig. 5A, C;). The MR signal contrast distinction observed between these basal ganglia structures matched well with the staining differences found in PV and ChAT sections (Figure 5B; the ChAT section is not shown). Different MRI parameters can visualize a clear separation (or lamina) between the putamen and pallidum or within the pallidum. The lateral medullary lamina (lml) and medial medullary lamina (mml) that separate pu from GPe, and GPe from GPi, respectively, are visible on the MTR and PV sections, but this separation is barely visible in T2W images (Fig. 5A-F). We found two distinct architectonic regions in GPi, separated by the lamina-like darkly-stained band in PV sections (red stars in E). This band is not found in MTR or T2W images. The lateral part of the GPi (GPi-L) revealed a mediolaterally oriented band of darkly-stained neuronal processes. In contrast, the medial part of the GPi (GPi-M) showed a dense network of honeycomb-like neuronal processes (Fig. 5E). In addition, all subregions of the globus pallidus (GPe, GPi-L, and GPi-M) reveal mosaic-like patterns with bright and dark contrasts in several MRI parameters. This architectonic pattern is most prominent in MTR and T2W images and can be comparable to the distinct neuropil compartments observed in matched PV-stained sections (Fig. 5D-F).

**Fig. 5.**
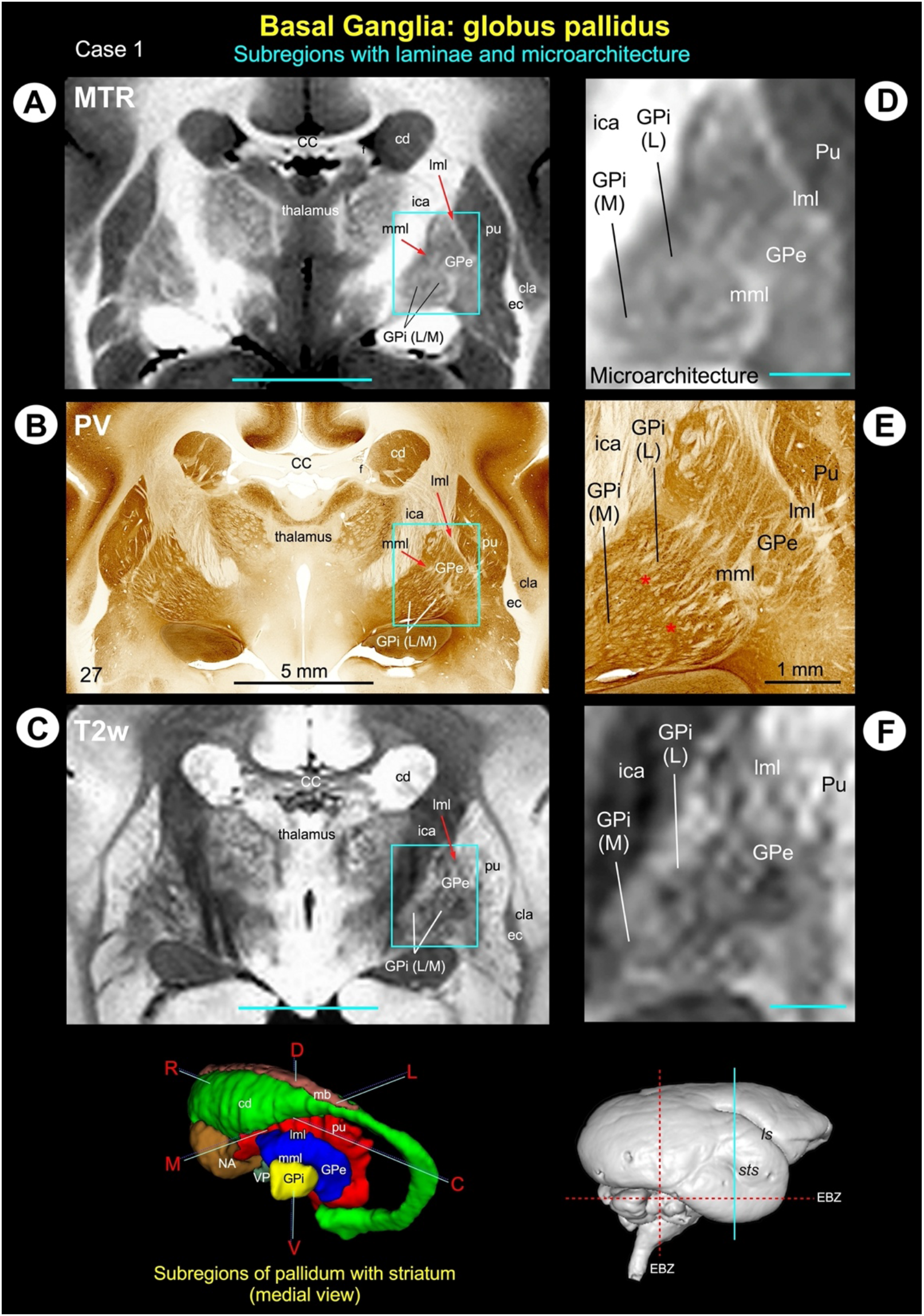
Pallidum. **(A-C)** illustrate the two subregions of the pallidum: globus pallidus external segment (GPe) and globus pallidus internal segment (GPi), medial to the putamen (pu) on the coronal MTR, and T2W images with matched histology section stained with parvalbumin (PV). **(D-F)** The zoomed-in view of the globus pallidus from A-C (green box) show the Pu, GPe, lateral, and medial subregions of the GPi (GPi-L and GPi-M, respectively) show the mosaic-like microarchitectural patterns in these subregions of the basal ganglia. The rostrocaudal extent of this coronal slice (green vertical line on the rendered brain image) and the spatial location and overall extent of the pallidum with other basal ganglia subregions in 3D are shown at the bottom. Note that both GPe and GPi are less hypointense or more hypointense than the laterally adjacent pu in MTR and T2W images, respectively, but significantly less hypointense than the surrounding white matter fiber tracts: the anterior limb of the internal capsule (ica) and the optic tract (ot) in T2W image (**A, C**). Red arrows in A-C indicate the location of the lateral medullary lamina (lml) between pu and GPe and the medial medullary lamina (mml) between GPe and GPi. These laminae are visible on the MTR and PV but barely visible in the T2w image. ***Abbreviations:*** CC-corpus callosum; cd-caudate nucleus; cla-claustrum; EBZ-ear bar zero; ec-external capsule; ica-internal capsule anterior limb; ot-optic tract. ***Sulci:*** ls-lateral sulcus; sts-superior temporal sulcus. ***Orientation on 3D:*** D-dorsal; V-ventral; R-rostral; C-caudal; M-medial; L-lateral. **Scale bar**: 5 mm (A-C); 1 mm applies to the magnified region in D-F.

#### Substantia nigra (Fig. 6)

The substantia nigra has been divided into two to four subregions by different studies based on the Nissl, myelin, AchE, tyrosine hydroxylase (TH), endogenous iron stains, or staining specific to dopamine cells in the midbrain (Arsenault et al., 1988; Francois et al., 1985; Jimenez-Castellanos and Graybiel, 1987). These subregions are the substantia nigra-pars reticulata (SNpr), pars compacta (SNpc), pars lateralis (SNpl), and pars mixta (SNpm). The marmoset SN was distinguished from the surrounding areas on different MRI parameters, but different subregions within SN were most strikingly discernible on the T2W images (Fig. 6A). We delineated all subregions with different contrast in this MRI parameter. The SNpc is readily identifiable as a compact mediolaterally oriented hyperintense band compared to the SNpl and SNpr subregions, as shown at the mid-level of the SN in Fig. 6A (right panel). The SNpl appears as a distinct hypointense region, overlapping with the adjacent less-hypointense SNpr and SNpm.

The spatial locations of these architectonic subregions matched well with the histological sections derived from the same brain specimen stained with SMI-32 or a different brain specimen stained with thionin (Fig. 6B, C). The SNpc is characterized by large, densely packed, and intensely stained cell bodies in thionin and darkly stained neurons with a network of neuronal processes in SMI-32 sections (Fig. 6B, C). This subregion has been known to contain dopaminergic cells (Arsenault et al., 1988). The SNpr is distinguished from SNpc by the presence of loosely packed small cell bodies in thionin but intensely-stained mosaic-like fiber bundles in SMI-32. The SNpl contains darkly-stained small to medium size cell bodies, which are more packed than the SNpr in thionin stain. This lateral region is a sparsely stained region with a network of thin neuronal processes in SMI-32. The SNpm is a narrow region with a mixture of different size neurons and lightly stained neuropil (Fig. 6B, C).

**Fig. 6.**
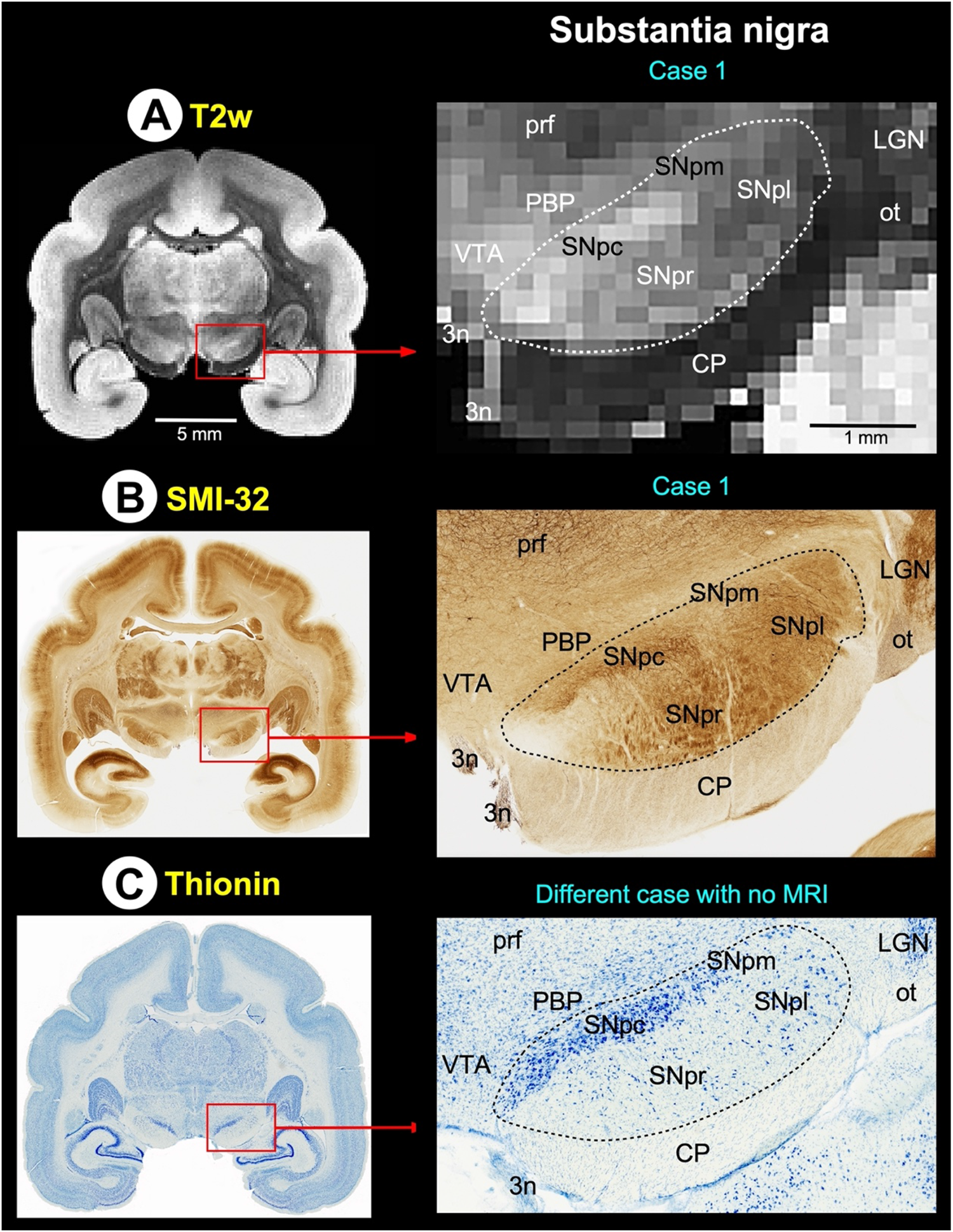
Substantia nigra. **(A-C)** Zoomed-in view of the red-boxed region shows three subregions of the substantia nigra: substantia nigra-pars compacta (SNpc), pars reticulata (SNpr), and pars lateralis (SNpl) with the adjacent ventral tegmental area (VTA), prerubral field (prf), and lateral geniculate nucleus (LGN) in coronal T2W, and matched histological sections stained with SMI-32 and thionin. Note that the thionin-stained section in C was obtained from a different marmoset monkey. ***Abbreviations:*** 3n-oculomotor nerve; CP-cerebral peduncle; ot-optic tract; PBP-parabrachial pigmented nucleus of the ventral tegmental area.

#### Red nucleus (Fig. 7)

The red nucleus (RN) extends for about 2.6 mm rostrocaudally and is divided into the rostral parvicellular (RNpc) region, which is contiguous with the caudal magnocellular (RNmc) region (Fig. 7). The location and overall extent of RN with reference to the basal ganglia subregions and a fiber bundle, the fasciculus retroflexus (fr), are illustrated in 3D (Fig. 7I). The contour of RNpc is less discrete from the surrounding gray and white matter structures in SMI-32, ChAT, and cresyl violet (CV) stained sections (Fig. 7B-D), and it contains small neurons scattered within the heterogeneously stained neuropil. The RNmc is sharply demarcated and contains large, intensely stained multipolar neurons identified in SMI-32 and CV-stained sections (Fig. 7F, H; inset). In contrast, this caudal subregion contains large but lightly-stained neurons and neuropil in ChAT stained section (Fig. 7G; inset). RNpc and RNmc are conspicuous regions with heterogeneous compartments of variable signal intensities, similar to the lateral part of the substantia nigra-SN in T2W images (Fig. 7A, E). The other subcortical regions surrounding the red nucleus: the oculomotor nuclei (3^rd^) and nerve (3n), the nucleus of Darkschewitsch (nd), the interstitial nucleus of Cajal (inc), and the interpeduncular nucleus (IPN), are delineated in both T2W and histology sections with different MR contrast and staining intensities of neuropil (Fig. 7).

**Fig. 7.**
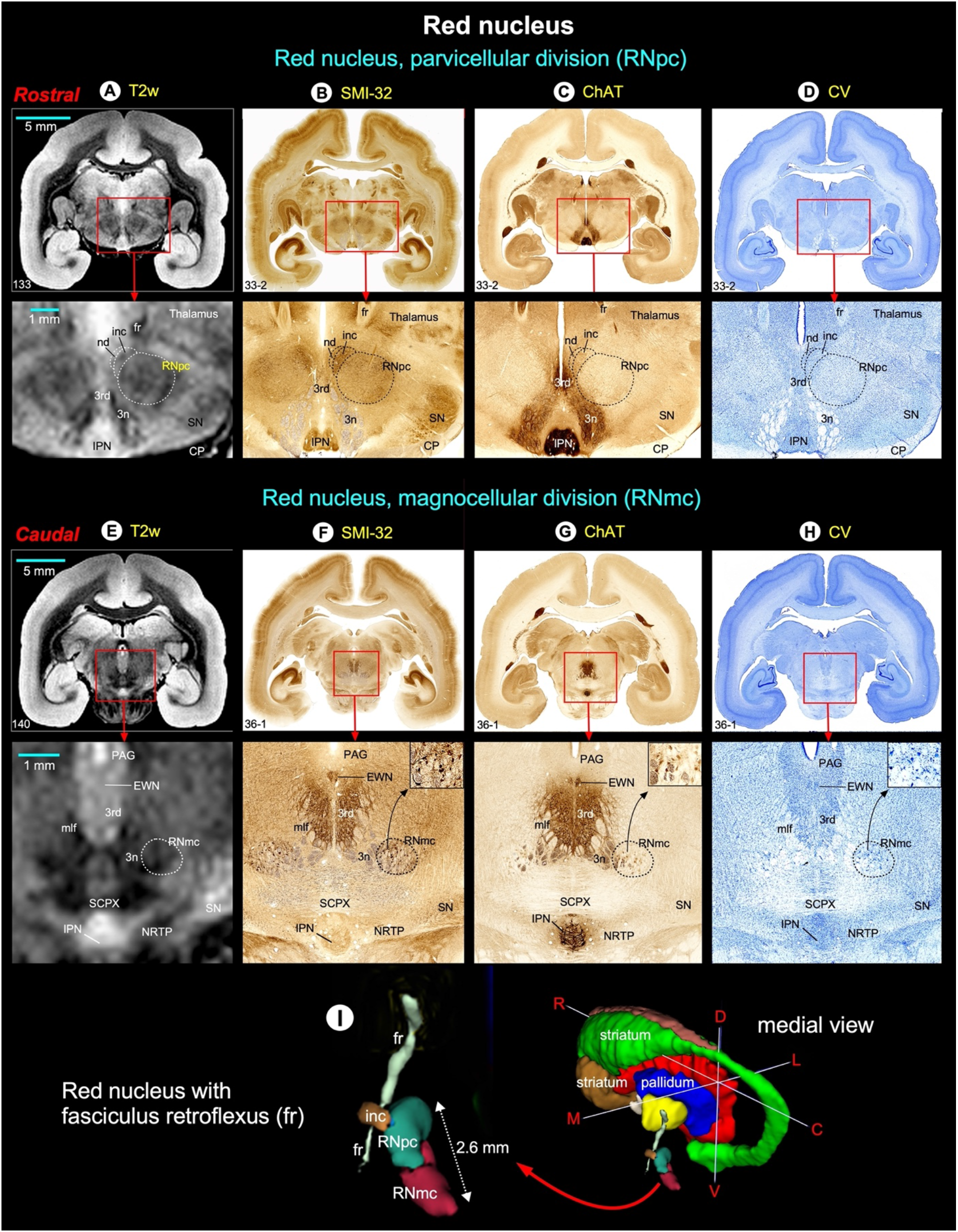
Red nucleus (Case 1). The zoomed-in view from the coronal midbrain slices (red boxes) shows the rostral parvicellular (RNpc; **A-D**) and the caudal magnocellular (RNmc; **E-H**) subregions of the red nucleus in relationship with the surrounding gray and white matter regions in T2W and matched histological sections. The dashed outlines on the T2W images indicate the corresponding location of the RNpc and RNmc derived from the matched histological sections. Insets in F-H indicate the magnified images of neurons and their processes in RNmc. We also identified other prominent gray and white matter regions surrounding the red nucleus at this midbrain level (see abbreviations below). **(I)** The 3D reconstruction shows the spatial location and rostrocaudal extent of the subregions of the red nucleus with reference to basal ganglia (striatum and pallidum) and the fiber bundle, the fasciculus retroflexus (fr). ***Abbreviations:*** 3rd-oculomotor nuclei; 3n-oculomotor nerve; CP-cerebral peduncle; EWN-Edinger Westphal nucleus; fr-fasciculus retroflexus; inc-interstitial nucleus of Cajal; IPN-interpeduncular nucleus; mlf-medial longitudinal fasciculus; nd-nucleus of Darkschewitsch; NRTP-nucleus reticularis tegmenti pontis; PAG-periaqueductal gray; SCPX-superior cerebellar peduncle decussation; SN-substantia nigra. Orientation: C-caudal; D-dorsal; L-lateral; M-medial; R-rostral; V-ventral.

### Fiber bundles associated with the basal ganglia (Fig. 8)

The fiber bundles associated with the striatum: Muratoff bundle (mb) on the dorsolateral part of the caudate nucleus (cd), anterior limb of the internal capsule (ica), and external capsule (ec) were identified on the track density (DEC-FOD) images (see Fig. 4). In addition, the pallidofugal axons that originate from different locations within the globus pallidus internal segment (GPi) form two fascicles, the ansa lenticularis (al) and lenticularis fasciculus (lf), and these fiber bundles course through Forel’s H fields (H, H1, and H2) on their way to the thalamus and brainstem (Baron et al., 2001; Lanciego et al., 2012; Neudorfer and Maarouf, 2018; Parent and Parent, 2004; Sidibe et al., 1997). The spatial locations of these fiber tracts with reference to Forel’s H field and gray matter structures such as the subthalamic nucleus (STN), zona inserta (zic), substantia nigra (SN), and thalamic nuclei were identified on coronal and sagittal DEC-FOD images and adjacent-matched SMI-32 and PV stained sections (Fig. 8).

The projections that originate from neurons located in the lateral and medial subregions of the GPi (Parent et al., 2001) form the ansa lenticularis (al) that course ventromedially around the posterior limb of the internal capsule (icp) and then bifurcates posteriorly to enter Forel’s H field (also known as the prerubral field; Fig. 8D). Similarly, axons from the lateral subregion of the GPi neurons course through the internal capsule to form the lenticular fasciculus (lf), located between STN and zona inserta (also called Forel’s field H2; Fig. 8D). Ultimately, the lenticular fasciculus merges with the ansa lenticularis at the level of field H of Forel to enter the thalamic nuclei through the thalamic fasciculus (H1 field of Forel) (Fig. 8D). The main thalamic targets of pallidum are the ventral anterior (VA), subregions of ventral lateral (VL, VLc, VLo), and the centromedian (cnMD) and intralaminar thalamic nuclei (Fig. 8A-D). The origin and direction of the fibers and projection targets illustrated in Figure 8D are based on previous anatomical tract tracing studies in macaque monkeys. For example, see (Parent and Parent, 2004).

**Fig. 8.**
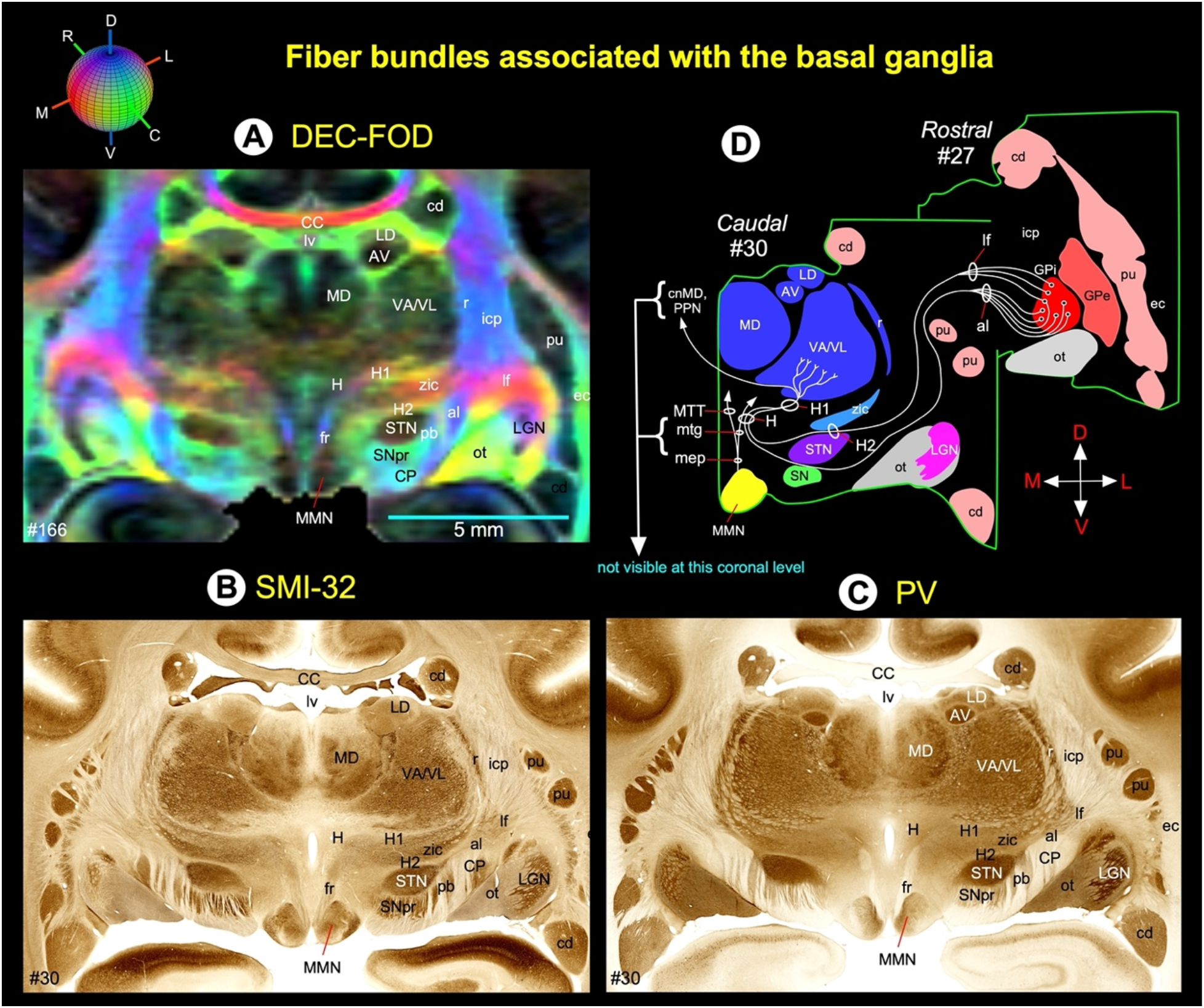
Fiber bundles linking basal ganglia and thalamus (**Case 1**). **(A-C)** Subregions of the basal ganglia, dorsal thalamus, and associated fiber bundles on coronal DEC-FOD image and corresponding histology section stained with SMI-32 and PV. Red, green, and blue in A indicate the direction of the fibers (anisotropy) along mediolateral, rostrocaudal, and dorsoventral directions, respectively. See the color-coded sphere with directions on the top. **(D)** The schematic diagram illustrates the projections from the GPi (internal segment of the globus pallidus) to different nuclei in the thalamus through ansa lenticularis (al), lenticular fasciculus (lf), and H, H1, and H2 fields of Forel. The direction of the fibers and projection targets is based on previous anatomical tracing studies in macaque monkeys. ***Abbreviations:*** ac-anterior commissure; al-ansa lenticularis; AV-anterior ventral nucleus; CC-corpus callosum; cd-caudate nucleus; cnMD-centromedian nucleus; CP-cerebral peduncle; ec-external capsule; f-fornix; fr-fasciculus retroflexus; GPe-globus pallidus external segment; GPi-globus pallidus internal segment; H, H1, H2-Fields of Forel; icp-internal capsule, posterior limb; LD-lateral dorsal nucleus; LGN-lateral geniculate nucleus; lf-lenticular fasciculus; lv-lateral ventricle; MD-medial dorsal nucleus of thalamus; mep-mammillary efferent pathway; MMN-medial mammillary nucleus; mtg-mammillotegmental tract; MTT-mammillothalamic tract; ot-optic tract; pb-pontine bundle; PPN-pedunculopontine tegmental nucleus; pu-putamen; r-reticular nucleus; SN-substantia nigra; SNpr-substantia nigra pars reticulata; STN-subthalamic nucleus; VA-ventral anterior nucleus; VL-ventral lateral nucleus; zic-zona incerta. ***Orientation:*** D-dorsal; V-ventral; R-rostral; C-caudal; M-medial; L-lateral.

### Comparison of basal ganglia subregions, red nucleus, and the deep cerebellar nuclei in marmoset and macaque monkeys (Fig. 9)

In T2W images, we noticed distinct MR signal intensity differences in some deep brain structures between marmoset and macaque monkeys. In particular, the subregions of the basal ganglia (ventral pallidum-VP, globus pallidus-GPe, GPi, and substantia nigra-SN) and the deep cerebellar nuclei (dentate-DN, interposed-AIN, PIN, and fastigial-FN) exhibited significantly more hypointense signal in the macaque than in the marmoset (Fig. 9A-J). We also confirmed this in another macaque and two marmoset T2W images (not shown here). Imaging studies in humans and non-human primates interpreted this hypointense signal to the site of preferential accumulation of iron-ferritin/hemosiderin (Bizzi et al., 1990; Drayer et al., 1986; Hardy et al., 2005; Manova et al., 2009). To confirm this finding, we stained matched histology sections from the same macaque and marmoset monkey brain specimens with Perl’s Prussian blue or iron stain (see method section).

#### Prussian blue stain in the Macaque monkey

Fig. 9K-N shows the matched histology sections from the macaque monkey brain stained with Prussian blue. The intense blue stained areas indicate the iron deposits, mainly confined to the basal ganglia subregions and the cerebellar nuclei (Fig. 9, right column). The spatial location of these intensely stained regions matched well with the decreased signal intensity found in these deep brain structures in T2W images (Fig. 9, F-H in the middle column). In the cerebellum, the dentate nucleus (DN) revealed more intense staining of neuropil with Prussian blue than the other deep cerebellar nuclei: the fastigial nucleus (FN) and the anterior and posterior subregions of interposed nuclei (AIN and PIN, respectively) (Fig. 9N). These differential staining of iron deposits matched well with the signal intensity differences observed between different deep cerebellar nuclei in T2W images (Fig. 9I).

The red nucleus (RN) subregions also showed different signal intensities in T2W images between these two non-human primate species. In contrast to the basal ganglia subregions, the marmoset’s parvicellular division of the RN (RNpc) exhibited more hypointense signal than in the macaque (Fig. 9C, H; red stars). The lack of strong hypointense contrast in the macaque RNpc matches well with the very weak Prussian blue staining in this subregion of the red nucleus (Fig. 9M, red stars). We also found other differences between these two species in the subcortical white matter. For example, the anterior limb of the internal capsule (ica), cerebral peduncle (CP), anterior commissure (AC), and the deep cerebellar white matter exhibited more hypointense signal in marmoset than in the macaque (Fig. 9; compare axial MRI slices in E and J).

#### Prussian blue stain in the Marmoset monkey (case 2)

As indicated in the method section, we did not have an additional set of histology sections from the 1^st^ marmoset brain (case 1) for the iron stain, but we have obtained a different perfusion fixed marmoset brain (case 2; Fig. 10) for both T2W MRI and histology (Prussian blue stain) to correlate the significantly less hypo-intense signal observed in the basal ganglia and the deep cerebellar nuclei (see above) with the intensity of iron deposit in this species.

Fig. 10A-D illustrates the basal ganglia subregions (VP, GPe/GPi, and SN) and the deep cerebellar nuclei in high-resolution T2W images (100 μm) from the 2^nd^ marmoset monkey, and Fig. 10E-H indicates the matched histology sections from the same monkey brain stained histologically with Prussian blue. The lightly or weakly stained blue areas on the right column indicate the iron deposits in these deep brain regions, including the adjacent white matter (e.g., the ica-anterior limb of the internal capsule). The spatial location of these weakly stained regions matched well with the increased signal intensity found in these deep brain structures in T2W images (Fig. 10, compare A-D with E-H). Based on the weak neuropil staining with Prussian blue and significantly less hypointense signals in T2W images (Fig. 10 left column), we conclude that these deep brain structures in the marmoset contain less iron than in the macaque monkey.

**Fig. 9.**
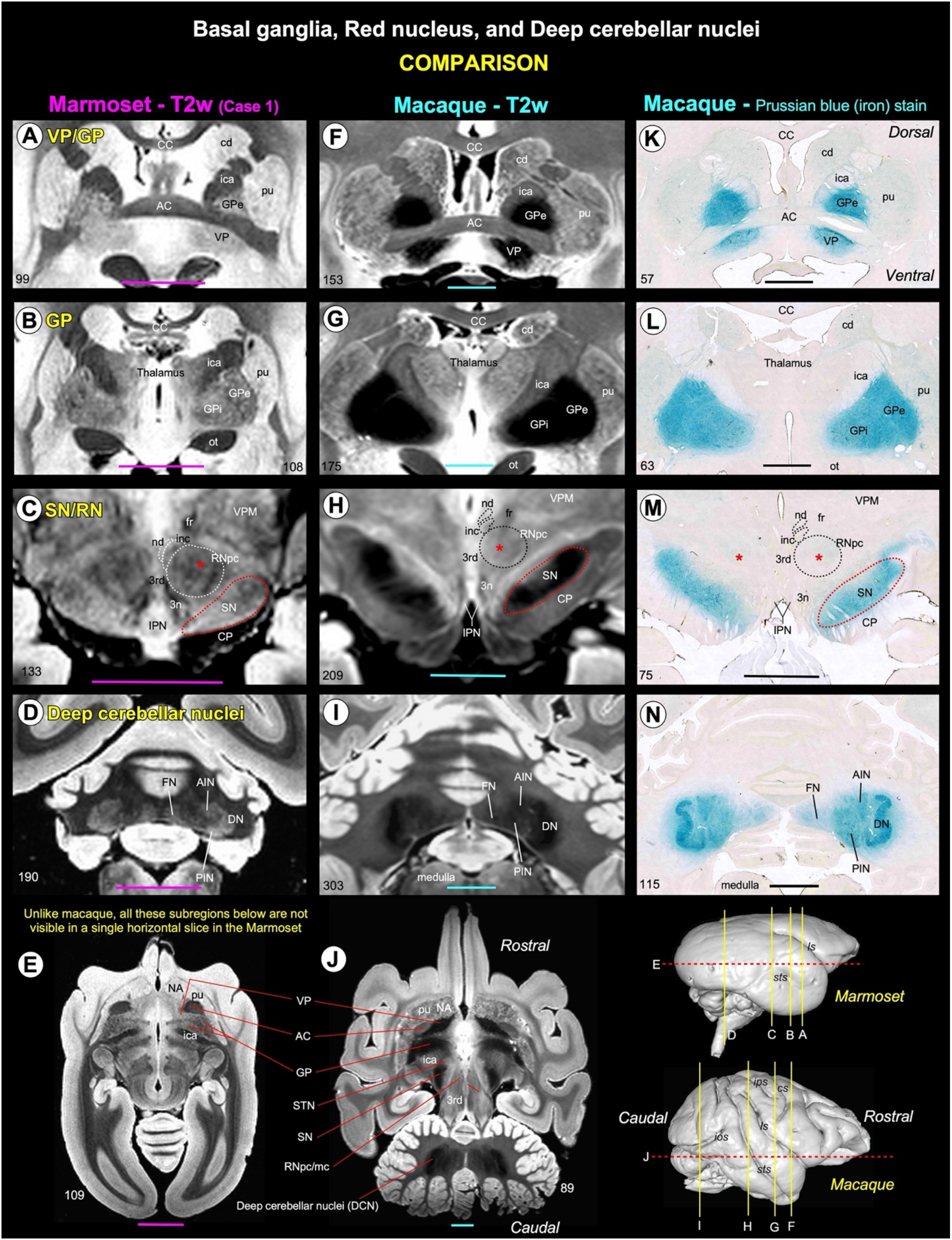
Comparison of subcortical regions in marmoset and macaque monkeys. **(A-E)** Subregions of the basal ganglia (caudate/putamen-cd/pu, ventral pallidum-VP, external and internal segments of the globus pallidus-GPe/GPi, substantia nigra-SN); parvicellular division of the red nucleus (RNpc); and the deep cerebellar nuclei (dentate nucleus-DN, anterior interposed nucleus-AIN, posterior interposed nucleus-PIN, fastigial nucleus-FN) in the marmoset **(case 1)** coronal and horizontal T2W images. **(F-J)** Similar subcortical regions in the macaque coronal and horizontal T2W images. **(K-N)** Matched histology sections with similar subcortical regions from the macaque monkey brain stained with Prussian blue for detecting intracellular ferric iron. Note that these basal ganglia subregions (except cd and pu) and the deep cerebellar nuclei stained intensely with Prussian blue that matched well with the strong hypointense signal found in these subregions, as revealed in T2W images. The rendered brain volumes of the marmoset and the macaque monkey at the bottom right show the location of coronal and horizontal T2W MRI slices A-J, as indicated by the solid yellow lines and red dashed lines, respectively, or the location of macaque coronal histology sections K-N, which are similar to F-I yellow lines. It is important to note that we do not have an additional set of histology sections from the marmoset brain (**case 1**) to confirm the presence of iron in these subregions by the histological method (but see Figure 10). ***Abbreviations:*** 3rd-oculomotor nucleus; 3n-oculomotor nerve; AC-anterior commissure; CC-corpus callosum; cd-caudate nucleus; CP-cerebral peduncle; DCN-deep cerebellar nuclei; ec-external capsule; fr-fasciculus retroflexus; GP-globus pallidus; GPe-globus pallidus external segment; GPi-globus pallidus internal segment; ica-internal capsule, anterior limb; inc-interstitial nucleus of Cajal; IPN-interpeduncular nucleus; NA-nucleus accumbens; nd-nucleus of Darkschewitsch; ot-optic tract; RN-red nucleus; RNmc-red nucleus, magnocellular division; RNpc-red nucleus, parvicellular division; STN-subthalamic nucleus; VP-ventral pallidum; VPM-ventral posterior medial nucleus; ***Sulci:*** cs-central sulcus; ios-inferior occipital sulcus; ips-intraparietal sulcus; ls-lateral sulcus; sts-superior temporal sulcus. ***Scale bars***: 5 mm.

**Fig. 10.**
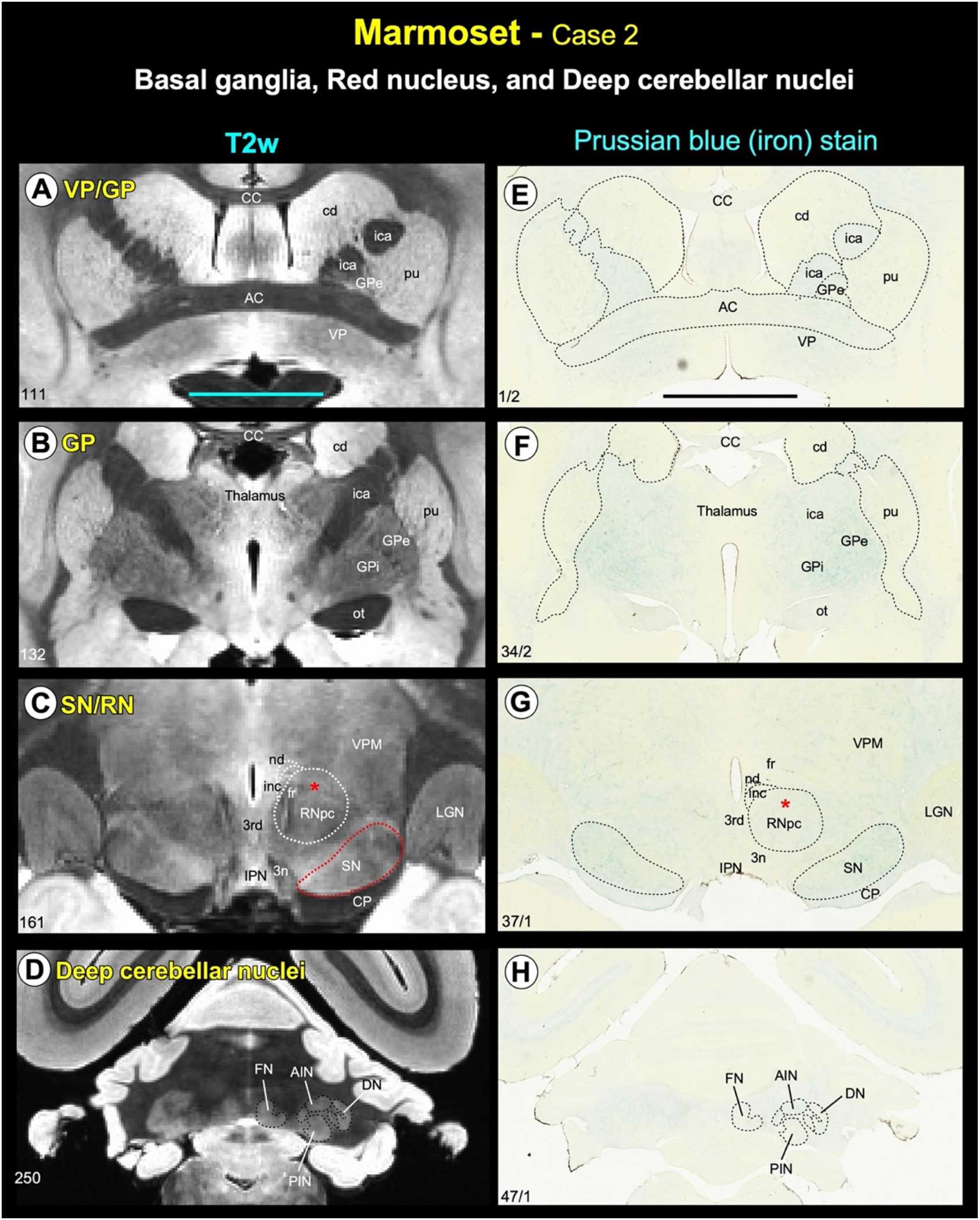
Comparison of subcortical regions in marmoset case 2. **(A-D)** Subregions of the basal ganglia (caudate/putamen-cd/pu, ventral pallidum-VP, external and internal segments of the globus pallidus-GPe/GPi, substantia nigra-SN); parvicellular division of the red nucleus (RNpc); and the deep cerebellar nuclei (dentate nucleus-DN, anterior interposed nucleus-AIN, posterior interposed nucleus-PIN, fastigial nucleus-FN) in the marmoset **(case 2)** coronal T2W images. **(E-H)** Matched histology sections with similar subcortical regions from the same marmoset monkey brain stained with Prussian blue for detecting intracellular ferric iron. Note that these basal ganglia subregions and the deep cerebellar nuclei stained weakly with Prussian blue that matched well with the significantly less hypointense signal found in these subregions, as revealed in T2W images. ***Abbreviations:*** 3^rd^-oculomotor nucleus; 3n-oculomotor nerve; AC-anterior commissure; CC-corpus callosum; cd-caudate nucleus; CP-cerebral peduncle; ec-external capsule; fr-fasciculus retroflexus; GP-globus pallidus; GPe-globus pallidus external segment; GPi-globus pallidus internal segment; ica-internal capsule, anterior limb; inc-the interstitial nucleus of Cajal; IPN-interpeduncular nucleus; LGN-lateral geniculate nucleus; nd-nucleus of Darkschewitsch; ot-optic tract; RN-red nucleus; SN-substantia nigra; VP-ventral pallidum; VPM-ventral posterior medial nucleus. **Scale bars**: 5 mm.

### Thalamus (Fig. 11)

The thalamic subregions in the marmoset are comparable to the macaque monkey (Olszewski, 1952). Thalamus is divided into the dorsal thalamus, epi-thalamus, and geniculate region (Jones, 1998) (Fig. 11). The dorsal thalamus is further divided into anterior, medial, lateral, intralaminar, and posterior groups and has significant roles in motor, emotion, memory, arousal, and other sensorimotor functions (Halassa and Kastner, 2017; Mitchell et al., 2014; Pergola et al., 2018).

The epi-thalamus is located in the posterior dorsal part of the diencephalon, and its principal gray matter structure is habenular nuclei, which play a pivotal role in reward processing, aversion, and motivation (Hikosaka et al., 2008; Roman et al., 2020). The geniculate region includes medial and lateral geniculate bodies and is an important relay nucleus in the auditory and visual pathways. We delineated these thalamic subregions in our MRI parameters, but the MAP-MRI parameter PA (propagator anisotropy) with DEC-FOD is particularly useful for identifying different thalamic nuclei and the surrounding fiber tracts of different orientations in the marmoset, as illustrated in Figure 11.

#### Dorsal Thalamus

##### Anterior and rostromedial groups

The anterior group of nuclei has a distinct pattern of connections with the hippocampal formation and other cortical areas, and it is thought to be involved in specific categories of learning and memory and spatial navigation (Aggleton et al., 2010; Jankowski et al., 2013; Shah et al., 2012). It is located in the dorsal part of the rostral thalamus and divided into the anterior dorsal (AD), anterior medial (AM), and anterior ventral (AV) nuclei. The AM, AV, and AD together form a distinct “V” shaped hypointense contrast band in PA/DEC-FOD image (Fig. 11A). This band is separated from the lateral group of thalamic nuclei (e.g., VAmc/VLc) ventrolaterally by a distinct hyperintense fiber bundle called the internal medullary lamina (iml; Fig. 11A). It continues with the dorsoventrally oriented mammillothalamic tract (MTT), which originates from the mammillary bodies. The caudal to the AV is the prominent lateral dorsal (LD) nucleus, which shows hypointense contrast similar to that of the AV (Fig. 11A, B). The hypointense contrast of the anterior thalamic nuclei in MRI corresponded well with the intensely stained neuropil in this thalamic region in the parvalbumin (PV) stained section (Fig. 11C). In contrast, this group of diencephalic nuclei is sparsely stained in SMI-32 (Fig. 11E).

The anterior group of nuclei is bordered dorsally by the medial group (paraventricular-Pa, parataenial-pt) and ventrally by the intralaminar group of nuclei (see below). Pa is located along the midline, adjacent to AM rostrally but adjacent to mediodorsal thalamic nuclei (MD) caudally (Fig. 11A, B). The pt is a small strip of hyperintense gray matter region on DEC-FOD with a limited rostrocaudal extent. The anterior group is also bordered dorsally by two distinct hyperintense fiber bundles, the stria medullaris (Sm) and fornix (f), which are easily delineated from the hypointense anterior group of nuclei (AM and AV) on the DEC-FOD image (Fig. 11A). The spatial location of these nuclei and fiber bundles is confirmed in matched histological sections, as shown in Figure 11C and E.

**Fig. 11.**
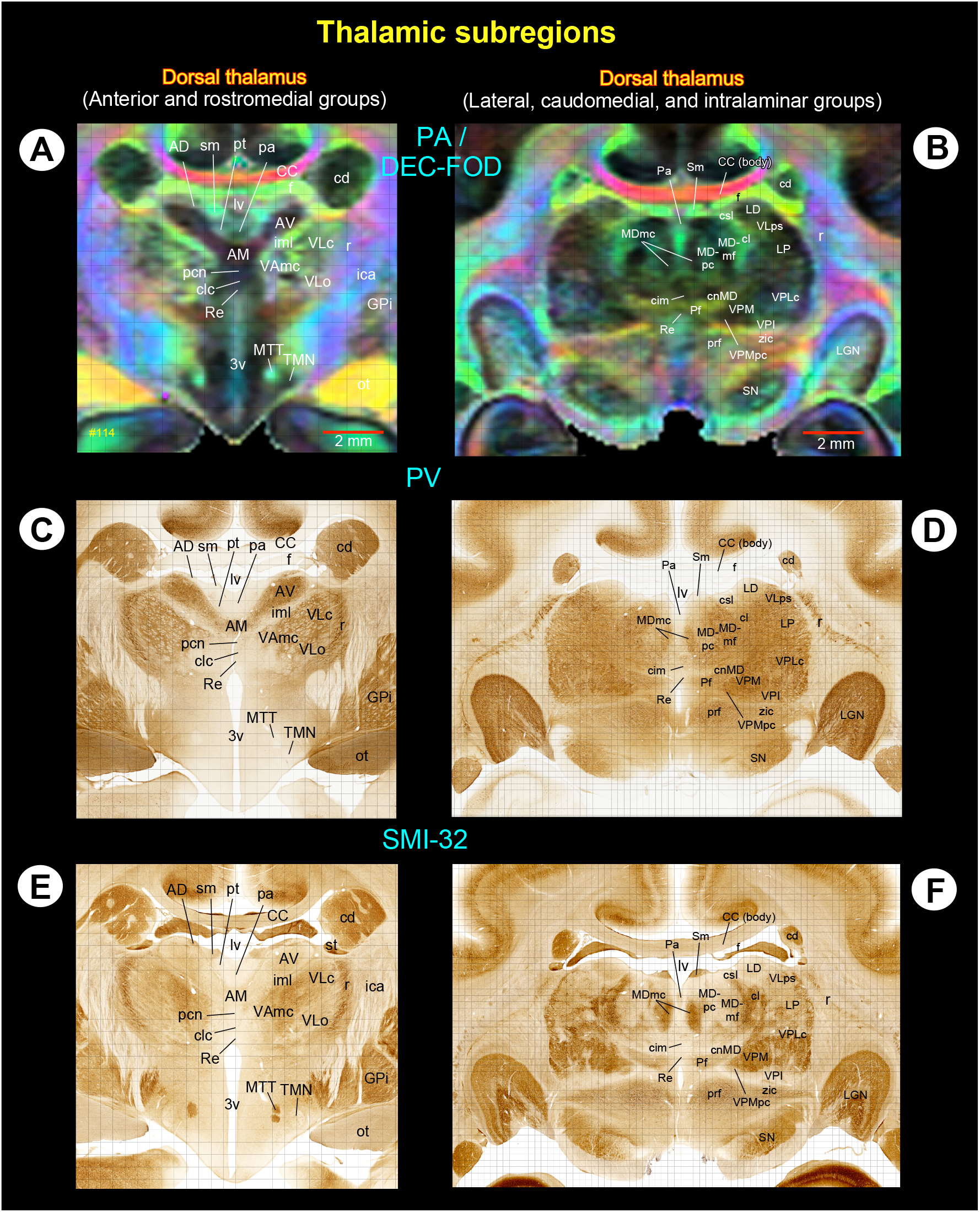
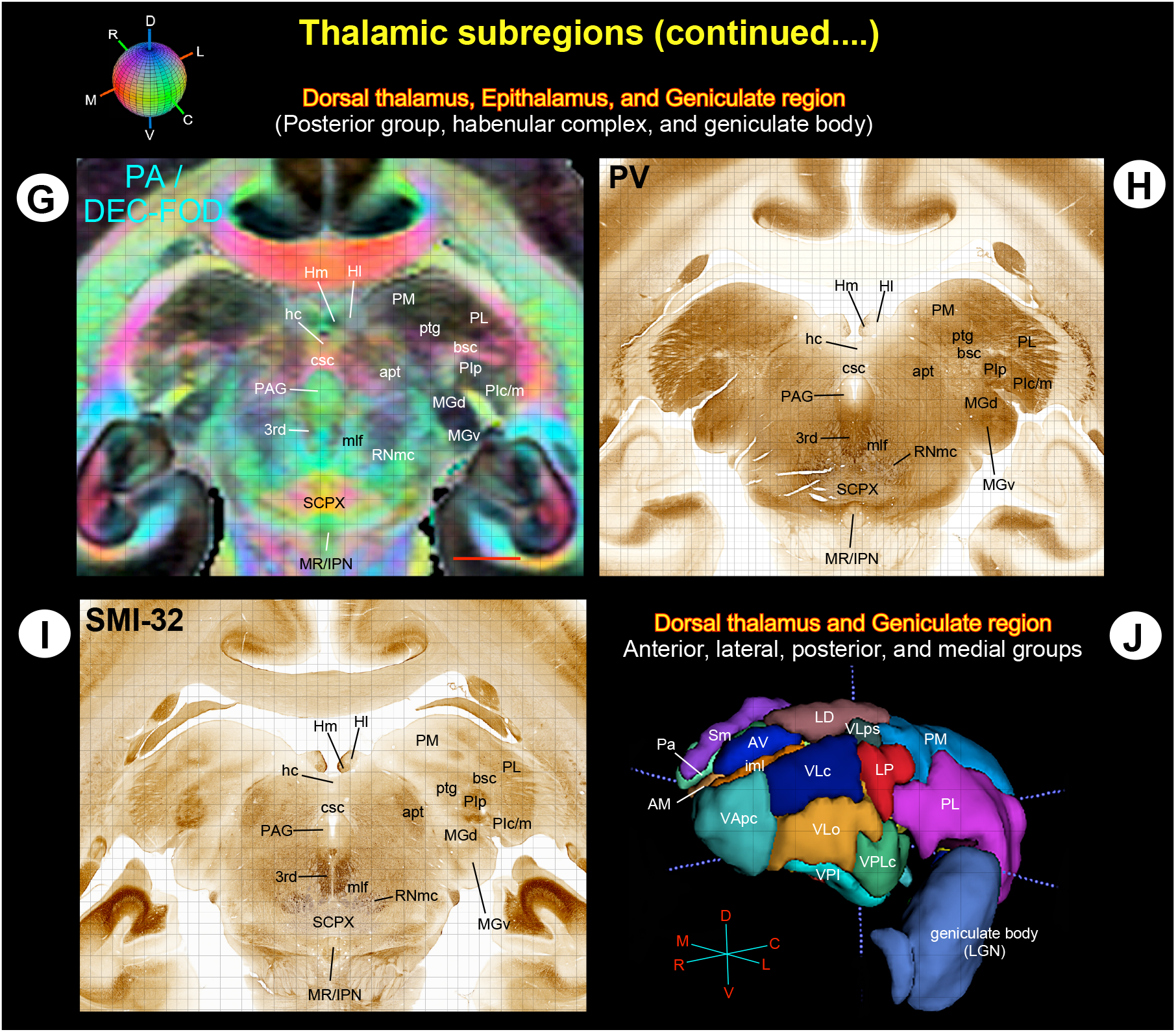
Thalamus (Case 1). **(A-I)** The MR signal intensity differences between dorsal thalamus (anterior, medial, intralaminar, and posterior groups), epithalamus (habenular complex), and geniculate region (geniculate body) in PA/DEC-FOD images, and the corresponding subregions in the histological sections stained with parvalbumin (PV), and SMI-32. **(J)** The spatial location and overall extent of the anterior, lateral, posterior, and some medial groups of thalamic nuclei in 3D reconstructed using ITK-SNAP. ***Abbreviations:*** 3^rd^-third cranial (oculomotor) nuclei; 3v-3^rd^ ventricle; bsc-brachium of superior colliculus; AD-anterior dorsal nucleus; AM-anterior medial nucleus; apt-anterior pretectal nucleus; AV-anterior ventral nucleus; bsc-brachium of superior colliculus; CC-corpus callosum; cd-caudate nucleus; cim-central intermediate nucleus; cl-central lateral nucleus; clc- central latocellular nucleus; cnMD-centromedian nucleus; csc-commissure of superior colliculus; csl-central superior lateral nucleus; f-fornix; GPi-globus pallidus, internal segment; hc-habenular commissure; Hl-lateral habenular nucleus; Hm-medial habenular nucleus; iml-internal medullary lamina; IPN-interpeduncular nucleus; LD-lateral dorsal nucleus; LGN-lateral geniculate nucleus; LP-lateral posterior nucleus; lv-lateral ventricle; MDmc-medial dorsal nucleus, magnocellular division; MDmf-medial dorsal nucleus, multiform division; MDpc-medial dorsal nucleus, parvicellular division; MGd-medial geniculate nucleus, dorsal division; MGv-medial geniculate nucleus, ventral division; mlf-medial longitudinal fasciculus; MR-median raphe; MTT-mammillothalamic tract; ot-optic tract; Pa-paraventricular nucleus; PAG-periaqueductal gray; pcn-paracentral nucleus; Pf-parafascicular nucleus; PI-inferior pulvinar; PL-lateral pulvinar; PM-medial pulvinar; pt-parataenial nucleus; prf-prerubral field; ptg-posterior thalamic group; r-reticular nucleus; Re-reunions nucleus; RNmc-red nucleus, magnocellular division; SCPX-superior cerebellar peduncle decussation; Sm-stria medullaris; SN-substantia nigra; TMN-tuberomammillary nucleus; VAmc-ventral anterior nucleus, magnocellular division; VApc-ventral anterior nucleus, parvicellular division; VLc-ventral lateral caudal nucleus; VLo-ventral lateral oral nucleus; VLps-ventral lateral postrema nucleus; VPI-ventral posterior inferior nucleus; VPLc-ventral posterior lateral caudal nucleus; VPLo-ventral posterior lateral oral nucleus; VPM-ventral posterior medial nucleus; VPMpc-ventral posterior medial nucleus, parvicellular division; zic-zona incerta. ***Orientation on 3D:*** D-dorsal; V-ventral; R-rostral; C-caudal; M-medial; L-lateral. Scale bars: 2 mm applies to A-I.

##### Lateral group

The lateral group is a large division of the thalamus with 11 subnuclei: ventral anterior (VApc, VAmc), ventral lateral (VLc, VLo, VLps), ventral posterior (VPLc, VPLo, VPM, VPMpc, VPI), and lateral posterior (LP) nuclei. The overall extent of these subregions is appreciated in 3D derived from tracing these nuclei in serial MRI sections using ITK-SNAP (Yushkevich et al., 2006) and confirmed with matched histological sections. (Fig. 11J). Anatomical tracing studies in non-human primates show that these nuclei represent the principal thalamic relays for inputs from the substantia nigra pars reticulata (SNpr), internal segment of the globus pallidus (GPi), deep cerebellar nuclei, and/or medial/trigeminal lemniscus (Asanuma et al., 1983; Kaas, 2012; Rouiller et al., 1994; Sakai et al., 1996).

The VAmc (ventral anterior, magnocellular division), VLo (ventral lateral oral), and VLc (ventral lateral caudal nucleus) are located in the rostral part of the lateral group. The VAmc and VLc exhibited more hypointense contrast than the VLo in DEC-FOD images (Fig. 11A, B). The VLc is continuous caudally with the LP, which shows a more hypointense signal than the VLc. Similarly, the rostrally located VLo exhibited a more hyperintense contrast than the caudally located contiguous area VPLc (Fig. 11A, D). The staining intensity of neuropil is also different in these regions in PV and SMI-32 stained sections (Fig. 11C, E).

The ventral posterior and lateral posterior nuclei dominate the caudal portion of the lateral group. Among this group, the VPM (ventral posterior medial), VPMpc (ventral posterior medial, parvicellular division), and LP (lateral posterior) revealed more hypointense contrast than the adjacent VPI (ventral posterior inferior) and VPLc (ventral posterior lateral caudal) in DEC-FOD. The VPI is the lightly stained region compared to other nuclei in this group in SMI-32 (Fig. 11F).

##### Caudomedial group

The prominent caudomedial group includes the mediodorsal thalamus with three subregions (MDmc, MDpc, and MDmf). It is thought to be involved in many cognitive tasks, such as decision-making, working memory, and cognitive control related to the prefrontal cortex (Ouhaz et al., 2018). In DEC-FOD, the MDmc (mediodorsal magnocellular) and MDmf (mediodorsal multiform) nuclei exhibited a hypointense contrast compared to MDpc (mediodorsal parvicellular), and this feature coincided with the dark-and lightly stained neuropil in these subregions, respectively, in PV stained section (Fig. 11D). In contrast, the MDmc is the most intensely stained region compared to the sparsely stained MDpc and MDmf in the SMI-32 (Fig. 11F).

##### Intralaminar groups

The intralaminar groups with ten nuclei (pcn, Clc, cdc, cim, cif, cl, Re, csl, cnMD, and Pf) are distributed rostrocaudally along the midline of the thalamus (Fig. 11). Due to the small size (except cnMD and Pf), the spatial location of these nuclei on MRI was challenging to distinguish from neighboring structures. These midline nuclei were marked on the MR images based on the spatial location and staining patterns observed in histologically stained sections. The cnMD (centromedian) and Pf (parafascicular) nuclei are visible as mediolaterally oriented bands, and the distinction between these subregions is less prominent in DEC-FOD and matched histology sections (Fig. 11B, D, F).

##### Posterior group

The posterior group is dominated by medial, lateral, and inferior pulvinar nuclei (PM, PL, and PI, respectively) and strongly connected with the visual cortical areas in the dorsal and ventral stream. The brachium of the superior colliculus, a prominent mediolaterally oriented fiber bundle, demarcates PM and PL from different subregions of PI and the medial geniculate body (MGB, part of the geniculate region, see below) (Fig. 11G). The PL and PI exhibited greater hypointense contrast relative to the medial part of the PM in DEC-FOD. This feature corresponded well with the dark and light neuropil staining in these subregions, respectively, in PV and SMI-32 stained sections (Fig. 11G-I).

#### Epi-thal3amus

The medial and lateral subregions of the habenular nuclei (Hm and Hl, respectively) and the associated fiber bundle, the habenular commissure (hc), form the principal gray and white matter structures of the epithalamus. The Hm is more hypointense than Hl in the PA/DEC-FOD image (Fig. 11G). The neuropil within the Hm is more intensely stained than in Hl in PV and SMI-32 (Fig. 11H, I).

#### Geniculate region

It comprises the lateral geniculate nucleus and the medial geniculate body (LGN and MGB, respectively). The contour of the LGN is easily distinguished from the surrounding regions in MR images, but the signal intensity differences between different laminae within LGN are less distinct except for the outer parvocellular layers 5 and 6, which exhibited more hypointense contrast than the magno (layers 1 and 2) and the inner parvocellular layers 3 and 4 (Fig. 11B). The MGB is divided into different subregions (e.g., MGd and MGv) and can be distinguished based on different signal intensities in MRI and staining patterns in histological sections (Fig. 11G-I).

### Brainstem (Figs. 12, 13, 14, and Suppl fig. 1)

The high-resolution MAP-MRI (PA with DEC-FOD) and T_2_w images are instrumental in delineating gray and white matter subregions in the different rostrocaudal extents of the brainstem (midbrain, pons, and medulla) and cerebellum. We delineated several sensory and motor nuclei and fiber tracts of different sizes and directions in the brainstem on three sagittal DEC-FOD images spaced 0.6 mm, 1.2 mm, and 2.4 mm from the midline (Fig. 12A-C). The direction of the fiber bundles was confirmed with fODFs (fiber orientation distribution function) as illustrated on the directionally encoded color images in A-C. We also delineated the selected brainstem nuclei in coronal high-resolution T2W images with 100 μm resolution from case 2, confirmed with matched histological sections stained with NeuN, SMI-32, and ChAT (Figs. 13 and 14). The architectonic subregions of the brainstem nuclei were identified with reference to the photographic atlas of the human brain (DeArmond et al., 1989) and Duvernoy’s atlas of the human brainstem and cerebellum (Naidich et al., 2009). We also used the marmoset brain atlas (Paxinos et al., 2012) to identify selected nuclei in the brainstem. The spatial location of some brainstem nuclei and fiber bundles in our marmoset MRI is comparable to macaques (Saleem et al., 2021). In addition to our multiple datasets from cases 1 and 2, we also used high-resolution sagittal images (80 μm resolution) from previously published and publicly shared diffusion MRI volumes of the marmoset (Liu et al., 2020) to delineate and confirm the location of the brainstem nuclei and fiber tracts (see Suppl. Fig. 1).

#### Midbrain

Three well-defined structures with low diffusion anisotropy are identified at the midbrain level on the DEC-FOD image, 0.6 mm lateral to the midline. These structures are the periaqueductal gray (PAG), oculomotor (3^rd^), and trochlear (4^th^) cranial nerve nuclei (Figs. 12A, D; the 4^th^ nucleus is not visible at this coronal level). The PAG is located ventromedial to the superior and inferior colliculi (SC and IC) and crossing fibers associated with the colliculi, the commissures of the superior and inferior colliculi (csc and cic, respectively). The 3^rd^ and 4^th^ cranial nerve nuclei are situated dorsal to the prominent rostrocaudally oriented medial longitudinal fasciculus (mlf) tract and mediolaterally oriented superior cerebellar peduncle decussation (SCPX) fibers (Fig. 12A, D). The midbrain reticular formation (MRF), a sizeable diffuse region with indistinct border and variable signal intensities, occupies the lateral part of the midbrain, as shown in the DEC-FOD images (Fig. 12B, D). The other major gray matter structure spanning the entire rostrocaudal extent of the midbrain is the pontine nuclei (PN). The PN is most strikingly delineated from rostrocaudally oriented corticospinal/corticobulbar tracts (CST/CBT or pyramidal tract-PT) and mediolaterally oriented superficial and deeper parts of the pontocerebellar fibers (pcf-s and pcf-d, respectively), as revealed by fODFs with DEC map (Figs. 12A-D; 12D). These subregions are less distinct and spatially not distinguishable from each other on a conventional T1W or T2W MRI (Fig. 16).

The other structures located rostral to the midbrain in the diencephalon region with variable signal intensities are the red nucleus (RNpc and RNmc), different subregions of the substantia nigra (SN), subthalamic nucleus (STN), zona incerta (zic), and associated fiber bundles H1 and H2, which are described in detail in the basal ganglia section (see Figs. 6-8). Ventral to RN is the ventral tegmental area (VTA), which exhibited low anisotropy in the DEC-FOD image (Fig. 12B). The other prominent fiber tracts with different orientations: the stria medullaris (Sm), habenular interpeduncular tract or the fasciculus retroflexus (THI/fr), postcommissural fornix (pocf), and the mammillothalamic tract (MTT) are visible on the sagittal slice spaced 1.2 mm from the midline (Fig. 12B). Sm innervates the habenular nucleus, and THI/fr forms the output of the habenular nucleus and projects to the midbrain and hindbrain monoaminergic nuclei (Roman et al., 2020). The pocf originates from the hippocampus and projects to the mammillary bodies, which in turn terminates on the anterior thalamic nuclei through MTT (Aggleton et al., 2010).

**Fig. 12.**
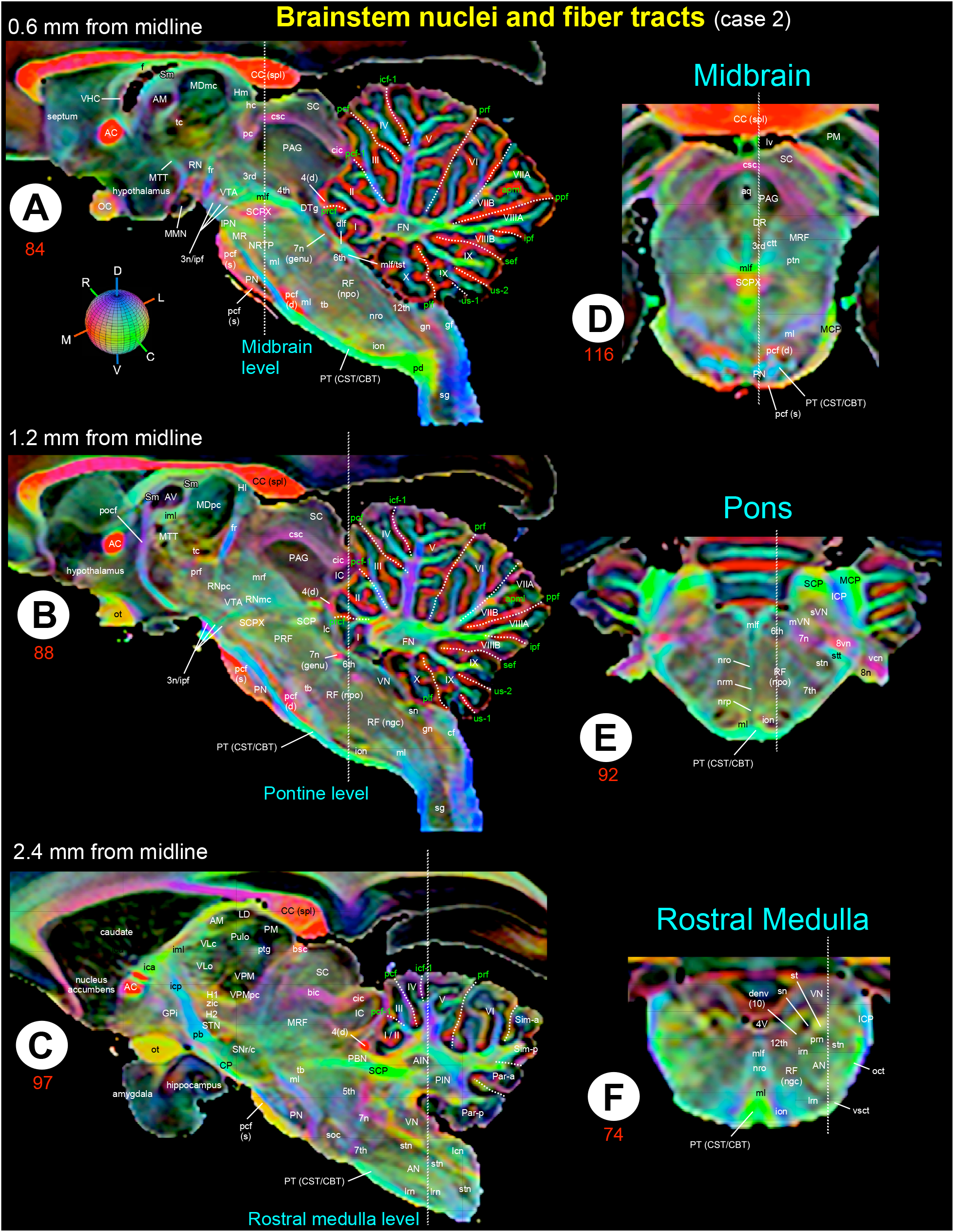
Brainstem and cerebellum. **(A-C)** Mediolateral extent of the brainstem (midbrain, pons, and medulla) with the cerebellum in three sagittal DEC-FOD images, spaced 0.6 mm, 1.2 mm, and 2.4 mm from the midline in case 2. ***Brainstem***: white or black letters indicate the gray matter regions or nuclei, and the major fiber tracts run in different directions in the brainstem (see the color-coded sphere with directions at the top in A). Red, green, and blue in these images indicate the anisotropy along mediolateral (ML), rostrocaudal (RC), and dorsoventral (DV) directions, respectively. Three dashed lines passing through the sagittal sections indicate the coronal sections at the midbrain, pons, and medulla level (D, E, F). The dashed lines on the coronal sections indicate the location of sagittal images illustrated on the left (A-C). Other gray and white matter subregions rostral to the brainstem (e.g., thalamus and basal ganglia) are also included. ***Cerebellum***: white letters indicate different cerebellar lobules, green letters show the names of cerebellar sulci and fissures, and white dashed lines show the sulci separating different lobules. ***Abbreviations for Brainstem and surrounding regions (A-C):*** 3^rd^-oculomotor nuclei; 3n-oculomotor nerve; 4^th^-trochlear nuclei; 4(d)-trochlear nerve decussation; 4V-4^th^ ventricle; 5^th^-trigeminal nuclei; 6^th^-abducent nuclei; 7^th^-facial nuclei; 7n-facial nerve; 7n (genu)-genu of the facial nerve; 8n-vestibulo-cochlear nerve; 8vn-vestibular nerve; 12^th^-hypoglossal nucleus; AC or ac-anterior commissure; AM-anterior medial nucleus; AN-ambiguous nucleus; aq-aqueduct; AV-anterior ventral nucleus; bic-brachium of inferior colliculus; bsc-brachium of the superior colliculus; CBT-corticobulbar tract; CC (spl)-splenium of corpus callosum; cf-cuneate fasciculus; cic-commissure of inferior colliculus; cn-cuneate nucleus; CP-cerebral peduncle; csc-commissure of superior colliculus; CST-corticospinal tract; ctt-central tegmental tract; denv (10)-dorsal efferent nucleus of vagus; dlf-dorsal longitudinal fasciculus; DR-dorsal raphe; dsct-dorsal spinocerebellar tract; DTg-dorsal tegmentum; dttt-dorsal trigemino thalamic tract; f-fornix; fr-fasciculus retroflexus; gf-gracile fasciculus; gn-gracile nucleus; GPi-globus pallidus internal segment; H1-H1 field of Forel; H2-H2 field of Forel; hc-habenular commissure; Hl-lateral habenular nucleus; Hm-medial habenular nucleus; IC-inferior colliculus; ica-anterior limb of the internal capsule; ICP-inferior cerebellar peduncle; icp-posterior limb of the internal capsule; iml-internal medullary lamina; ion-inferior olivary nucleus; ipf-inter peduncular fossa; IPN-interpeduncular nucleus; irn-intermediate reticular nucleus; lc-locus coeruleus; lcn-lateral cuneate nucleus; LD-lateral dorsal nucleus; lrn: lateral reticular nucleus; lv-lateral ventricle; MCP-middle cerebellar peduncle; MDmc-medial dorsal nucleus, magnocellular division; MDpc-medial dorsal nucleus, parvicellular division; ml-medial lemniscus; mlf-medial longitudinal fasciculus; MMN-medial mammillary nucleus; MR-median raphe; mrf-mesencephalic reticular formation; MRF-midbrain reticular formation; MTT-mammillothalamic tract; mVN-medial vestibular nucleus; nrm-nucleus raphe magnus; nro-nucleus raphe obscurus; nrp-nucleus raphe pallidus; NRTP-nucleus reticularis tegmenti pontis; OC-optic chiasm; oct-olivocerebellar tract; ot-optic tract; PAG-periaqueductal gray; pb-pontine bundle; PBN-parabrachial nucleus; pc-posterior commissure; pcf (d)-pontocerebellar fiber, deeper part; pcf (s)-pontocerebellar fibers, superficial part; pd-pyramidal decussation; PM-medial pulvinar; PN-pontine nuclei; pocf-postcommissural fornix; PRF-pontine reticular formation; prf-prerubral field; prn- parvicellular reticular nucleus; PT-pyramidal tract; ptg-posterior thalamic group; ptn-pedunculotegmental nucleus; Pulo-pulvinar oralis nucleus; RF (ngc)-reticular formation, nucleus gigantocellularis; RF (npo)-reticular formation, nucleus pontis centralis oralis; RN-red nucleus; RNmc-red nucleus, magnocellular division; RNpc-red nucleus, parvicelluar division; SC-superior colliculus; SCP-superior cerebellar peduncle; SCPX-superior cerebellar peduncle decussation; sg-substantia gelatinosa; Sm-stria medullaris; sn-solitary nucleus; SNr/c-substantia nigra pars reticulata/compacta; soc-superior olivary complex; st-solitary tract; stn-spinal trigeminal nucleus; STN-subthalamic nucleus; stt-spinal trigeminal tract; sVN-spinal vestibular nucleus; tb-trapezoid body; tc-thalamic commissure; vcn-ventral cochlear nucleus; VHC-ventral hippocampal commissure; VLc-ventral lateral caudal nucleus; VLo-ventral lateral oral nucleus; VN-vestibular nuclei; vsct-ventral spinocerebellar tract; VPM-ventral posterior medial nucleus; VPMpc-ventral posterior medial nucleus, parvicellular division; VTA-ventral tegmental area; zic-zona incerta. ***Abbreviations for cerebellar regions (A-C):*** I to X-cerebellar lobules; AIN-anterior interposed nucleus; FN-fastigial nucleus; Par_a-paramedian lobule anterior part; Par_p-paramedian lobule posterior part; PIN-posterior interposed nucleus; Sim_a-anterior part of the simple lobule; Sim_p-posterior part of the simple lobule. ***Cerebellar sulci/fissures (A-C):*** apml-ansoparamedian lobule; icf1-intraculminate fissure 1; ipf-intrapyramidal fissure; pcf-preculminate fissure; pcf1-preculminate fissure 1; plf-posterolateral fissure; ppf-prepyramidal fissure; prcf-precentral fissure; prf-primary fissure; sef-secunda fissure; us1-uvular sulcus 1; us2-uvular sulcus 2.

#### Pons

Six major cranial nerve nuclei (trigeminal-5^th^, abducens-6^th^, facial-7^th^, vestibular-VN, spinal trigeminal-stn, and superior olivary complex-soc) with low to moderate diffusion anisotropy are distinguished in the different rostrocaudal and mediolateral extent of the pons on sagittal and coronal PA/DEC-FOD images (Fig. 12A-C). Our high-dimensional coronal T2W images with an isotropic resolution of 100 µm from case 2 revealed several gray (nuclei) and white matter regions with variable signal intensities in the pontine regions (Fig. 13A). These are the cranial nerve nuclei, as indicated above, pontine reticular formation (RF-npo), raphe nuclei (nro, nrm, and nrp), cerebellar peduncles (SCP, MCP, ICP), corticospinal/bulbar tract (CST/CBT), medial lemniscus (ml), facial nerve (7n), and vestibulocochlear nerve (8n). The spatial locations of these regions are confirmed and matched well with the histological sections from the same brain specimen stained with NeuN, ChAT, and SMI-32 (Fig. 13B-D).

The high-resolution T2W images also revealed subregions with distinct signal intensities within the cranial nerve nuclei. For example, the lateral (L) subregion of the 7^th^ (facial) nucleus exhibited significantly more hyperintense signals than the other subregions within this nucleus (Fig. 13A, I; blue square). This feature was also confirmed in different marmoset T2w images. The hyper-intense L region correlated well with the densely packed and intensely stained neurons in this subregion in NeuN and ChAT stained sections (Fig. 13J, K) but not SMI-32 (Fig. 13L). The 6^th^ (abducens) nucleus is located close to the midline and immediately ventral to the crossing fibers of the 7^th^ nerve, which forms the genu of the facial nerve or the facial colliculus-fc (Fig. 13A, E). Three subregions of the 6^th^ cranial nerve nuclei can be identified with different signal intensities in T2W images: the hyperintense ventrolateral (VL) and dorsomedial (DM) groups and the hypointense medial (M) group, as illustrated in Fig. 13A, E; red square. The hyperintense VL and DM subregions corresponded well with the densely packed neurons, whereas the hypointense M region correlated with the sparsely distributed neurons in the 6^th^ nucleus in NeuN, ChAT, and SMI-32 stained sections (Fig. 13F-H). Among these identified cranial nerve nuclei in the pons, the 7^th^ (facial) is the most intensely stained region in the SMI-32 stained section (Fig. 13D, L).

**Fig. 13.**
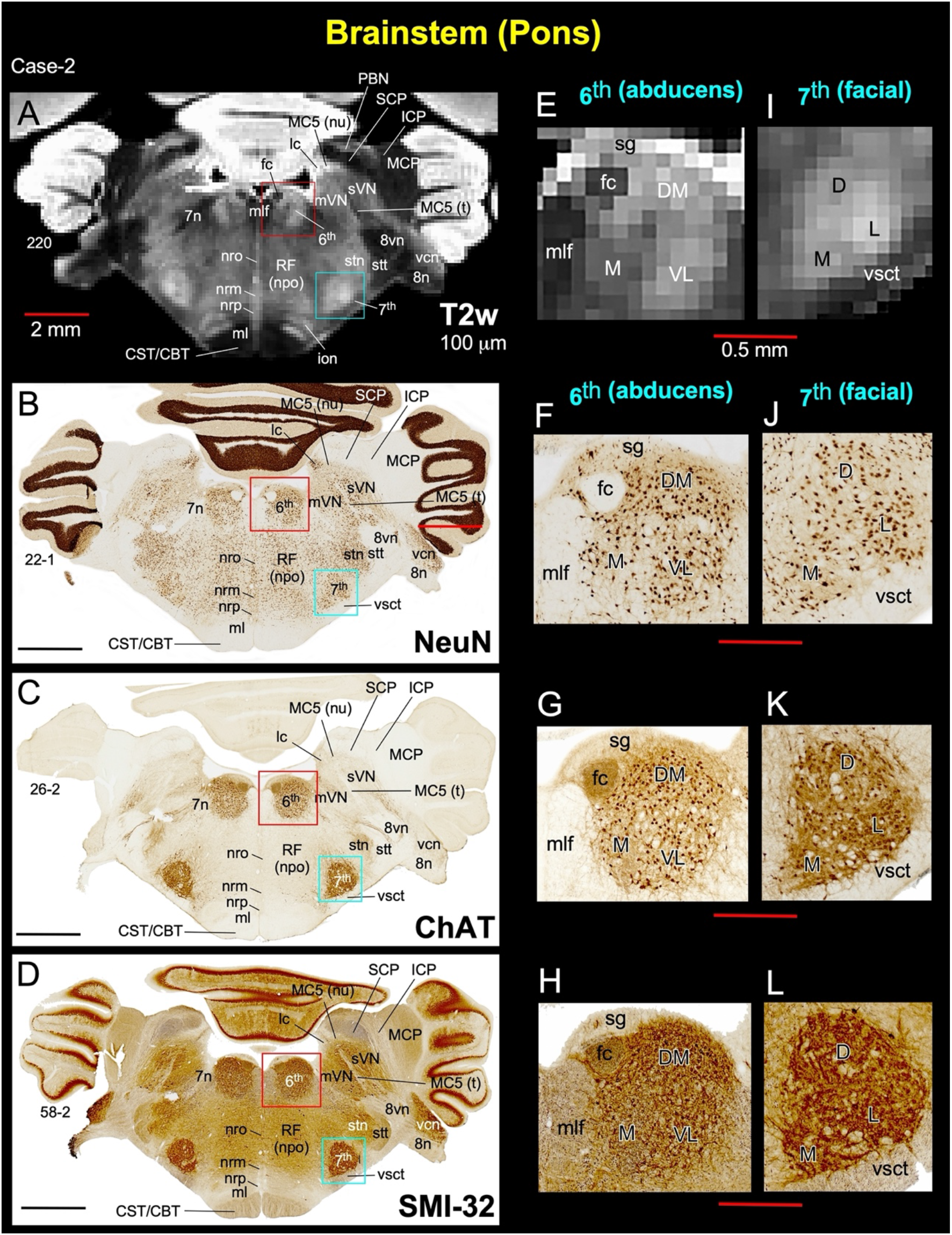
Brainstem (Pons). **(A-D)** The spatial location of different nuclei and fiber tracts at the pontine level in coronal T2W image (100 μm) and histology sections stained with NeuN, ChAT, and SMI-32. (**E-H**) The zoomed-in view of the 6^th^ (abducens) and 7^th^ (facial) cranial nerve nuclei indicated by the red and blue squares, respectively, on the images (left column), shows the subregions with different MR contrast and neuropil staining within these nuclei. ***Abbreviations:*** 6^th^-abducent nuclei; 7^th^-facial nuclei; 7n-facial nerve; 8n-vestibulocochlear nerve; 8vn-vestibular nerve; CBT-corticobulbar tract; CST-corticospinal tract; D-dorsal; DM-dorsomedial; fc-facial colliculus; ICP-inferior cerebellar peduncle; ion-inferior olivary nucleus; L-lateral; lc-locus coeruleus; M-medial; MC5 (nu)-mesencephalic trigeminal nucleus; MC5 (t)-mesencephalic trigeminal tract; MCP-middle cerebellar peduncle; ml-medial lemniscus; mlf-medial longitudinal fasciculus; mVN-medial vestibular nucleus; nrm-nucleus raphe magnus; nro-nucleus raphe obscurus; nrp-nucleus raphe pallidus; PBN-parabrachial nucleus; RF (npo)-reticular formation, nucleus pontis centralis oralis; SCP-superior cerebellar peduncle; stn-spinal trigeminal nucleus; stt-spinal trigeminal tract; sVN-superior vestibular nucleus; vcn-ventral cochlear nucleus; VL-ventrolateral; vsct-ventral spinocerebellar tract. Scale bar: 2 mm (A-D); 0.5 mm (E-L).

#### Medulla

Our high-resolution coronal T2W images at the level of the rostral medulla revealed several patchy gray matter regions with variable signal intensities (Fig. 14A, D). These are the nucleus prepositus (np), hypoglossal nucleus (12^th^), a dorsal motor nucleus of the vagus (denv-10), solitary nucleus (sn), cuneate nucleus (cn), medial and spinal vestibular nuclei (mVN and spVN, respectively), spinal trigeminal nucleus (stn), and the ambiguous nucleus (AN). The other gray matter structures include the subregions of the reticular formation: the nucleus gigantocellularis (RF-ngc) and lateral, intermediate, and parvicellular subregions of the reticular nucleus (lrn, irn, and prn, respectively). The spatial locations of these architectonic subregions matched well with the histological sections derived from the same brain specimen stained with NeuN and ChAT (Fig. 14B-F). Among the identified gray matter nuclei in the pons, the denv-10 (dorsal motor nucleus of the vagus) exhibited a strong hyperintense signal followed by the 12^th^ (hypoglossal) and np (nucleus prepositus) which showed a slightly less hyperintense signal (Fig. 14D). The corresponding and matched histology section stained with ChAT revealed more intensely stained neurons and neuropil in these three nuclear groups than other gray matter structures at this pontine level (Fig. 14F). In contrast, the denv (10^th^) and np are less prominent with sparsely stained neuronal cell bodies in compared to 12^th^ and rest of the gray matter structures in the NeuN stained section (Fig. 14E).

The rostroventral and medial part of the medulla is dominated by the well-defined hypointense inferior olivary nucleus (ion), which is strikingly delineated from ventrally adjacent pyramidal tract (CST/CBT) and dorsally located reticular formation (RF-ngc) in DEC-FOD images (Figs. 12B-F). The crossing fibers of the motor pathways: the pyramidal decussation (pd) dominates the caudal part of the medulla and exhibits higher anisotropy than the surrounding gray matter structure, the substantia gelatinosa (sg), as shown in Fig. 12A. The ambiguous nucleus (AN) is visible as a rostrocaudally extended hypointense region on the lateral part of the medulla. It can be demarcated from the surrounding hyperintense white matter tracts (Fig. 12C).

**Fig. 14.**
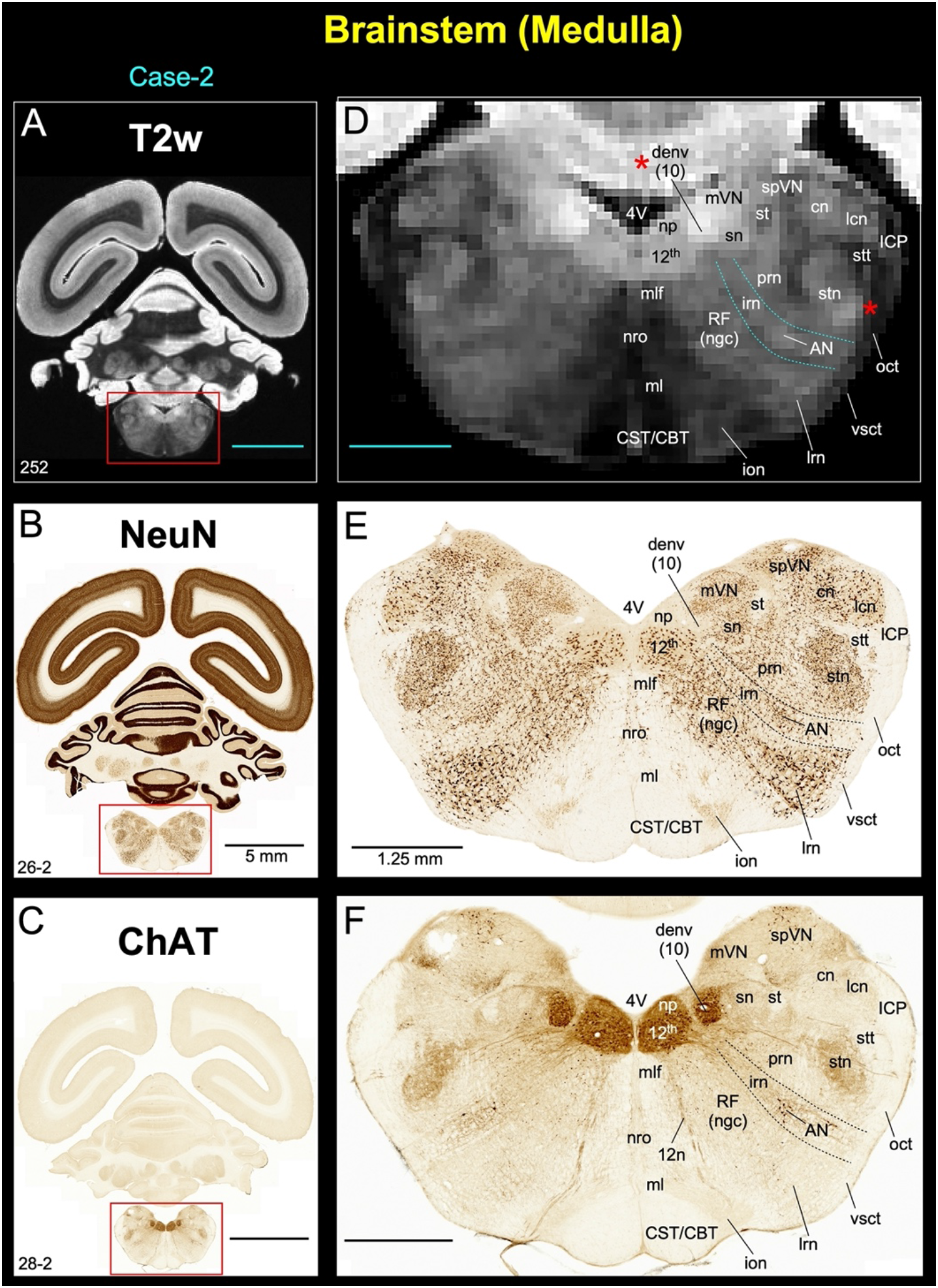
Brainstem (Medulla). **(A-C)** The spatial location of different nuclei and fiber tracts at the level of the rostral medulla in coronal T2W image (100 μm) and matched histology sections stained with NeuN and ChAT. Note the corresponding location of nuclei in both MRI and histology with variable MR and staining contrast. Also note the hyper-intense contrast of the dorsal motor nucleus of the vagus (denv-10) and intense staining of neurons in this nucleus, including hypoglossal (12^th^) and nucleus prepositus (np) in ChAT stained section **(D, F)**. ***Abbreviations:*** 12^th^-Hypoglossal nucleus; 4V-fourth ventricle; AN-ambiguous nucleus; cn-cuneate nucleus; CBT-corticobulbar tract; CST-corticospinal tract; denv (10)-dorsal motor nucleus of vagus; ICP-inferior cerebellar peduncle; ion-inferior olivary nucleus; irn-intermediate reticular nucleus; lcn-lateral cuneate nucleus; lrn-lateral reticular nucleus; ml-medial lemniscus; mlf-medial longitudinal fasciculus; mVN-medial vestibular nucleus; np-nucleus prepositus; nro-nucleus raphe obscurus; oct-olivocerebellar tract; prn-parvicelluar reticular nucleus; RF (ngc)-reticular formation, nucleus gigantocellularis; sn-solitary nucleus; spVN-spinal vestibular nucleus; st-solitary tract; stn-spinal trigeminal nucleus; stt-spinal trigeminal tract; vsct-ventral spinocerebellar tract. Scale bar: 5 mm (A-C); 1.25 mm (D-F).

### Cerebellum (Fig. 12 and Suppl Fig. 1)

Although unrelated to the present study, the cerebellar cortex’s unique foliation and compartmental organization prompted us to identify and segment this non-subcortical region in our high-resolution MR images. Similar to the macaque (Saleem et al., 2021), we identified the spatial location of 10 lobules (I-X), paramedian lobules (Par), simple lobule (Sim), ansiform lobules Crus I and II, flocculus (Fl), and paraflocculus (PFl) with reference to 11 landmarks (nine fissures and two sulci) through the rostrocaudal and mediolateral extent of the cerebellum on the sagittal MR images (Fig. 12A-C). The nomenclature and abbreviations used to identify these lobules and landmarks are according to (Larsell, 1953) but with some modifications to include the simplified version of the abbreviations for the fissures and sulci (see Fig. 12 legend). The cerebellar cortex lobules and associated fiber bundles’ orientation exhibited similar signal intensities in the different mediolateral extent of the cerebellum on DEC-FOD (Fig. 12A-C) and other MRI parameters.

The deep cerebellar nuclei (DCN), the sole output of the cerebellum, are vital structures of the cortico-cerebellar circuitry. In marmosets, four DCN embedded within the cerebellar white matter on each hemisphere were distinguished: the dentate nucleus (DN), anterior and posterior interposed nuclei (AIN and PIN), and the fastigial nucleus (FN). The AIN and PIN can be comparable to the emboliform and globose nuclei in humans (Tellmann et al., 2015). In T2W images, these four subregions of the DCN in the marmoset show more hyperintense signals than in the macaque monkey, which suggests the low iron content. For more details on the comparison, see figures 9 and 10.

### Amygdala (Fig. 15)

The nuclear subdivisions of the amygdaloid complex in the marmoset (Araujo Gois Morais et al., 2019) closely resemble those of macaque monkeys (Price et al., 1987; (Amaral and Bassett, 1989; Pitkanen and Amaral, 1998). The deep nuclei of the amygdala are divided into lateral (L), basal (B), accessory basal (AB), and paralaminar (PL’) nuclei. The lateral nucleus is further divided into four subregions (dorsal-Ld, lateral-Ll, medial-Lm, and ventral-Lv), defined based on cell size and packing density in Nissl and the intensity of neuropil staining in AchE (Araujo Gois Morais et al., 2019). We identified similar subregions in ChAT and NeuN stained sections.

Among these subregions, the Lv has weakly stained ChAT-positive fibers, and Ll has densely packed cell bodies in NeuN (Fig. 15C-D). The basal nucleus is divided into magnocellular, intermediate, and parvicellular subregions (Bmc, Bi, and Bpc, respectively). The Bmc and Bi stand out in ChAT with intense neuropil labeling and in NeuN with large to medium size cells compared to the Bpc (Fig. 15C-D). The accessory basal nucleus is delineated into the magnocellular (ABmc), parvicellular (ABpc), and ventromedial (ABvm) regions. The ABmc showed an intensely stained neuropil compared to ABpc and ABvm in ChAT (Fig. 15C). The ABmc held the largest cells, but the ABvm revealed more packed cells in NeuN (Fig. 15D).

Unlike the macaque monkey, the paralaminar nucleus (PL’) in the marmoset is a thick band of densely packed neurons located ventral to the other deep nuclei and dorsal to the entorhinal cortex (Fig. 15C-D). We delineated the histologically identified deep nuclei (L, B, and AB) in MRI images. In particular, these nuclei exhibited hypo- or hyperintense contrast in MAP-MRI (DEC-FOD) and T2W images, respectively, but the border between many of the subregions within these deep nuclei (see above) is less prominent in both MR images (Fig. 15A-B). We identified the borders on the MRI based on the contour of the nuclei derived from the matched ChAT sections.

The other subdivisions of the amygdala include the medial nucleus (ME), anterior and posterior cortical nuclei (Coa and Cop, respectively), medial and lateral subregions of the central nucleus (CEm and CEL), anterior amygdaloid area (AAA), and amygdalohippocampal area (AHA). The central nucleus is the prominent structure located dorsal to the basal nuclei, and it stands out in the ChAT section with intensely labeled neuropil (Fig. 15C). The ME is the sparsely stained region in ChAT among the identified amygdaloid nuclei. The central (CEm and CEL) nucleus exhibited a hypointense signal, whereas the medial-ME nucleus showed hyperintense contrast in DEC-FOD. The contrast found in these nuclei in DEC-FOD is the opposite of the signal found in T2W images (Fig. 15A-B).

**Fig. 15.**
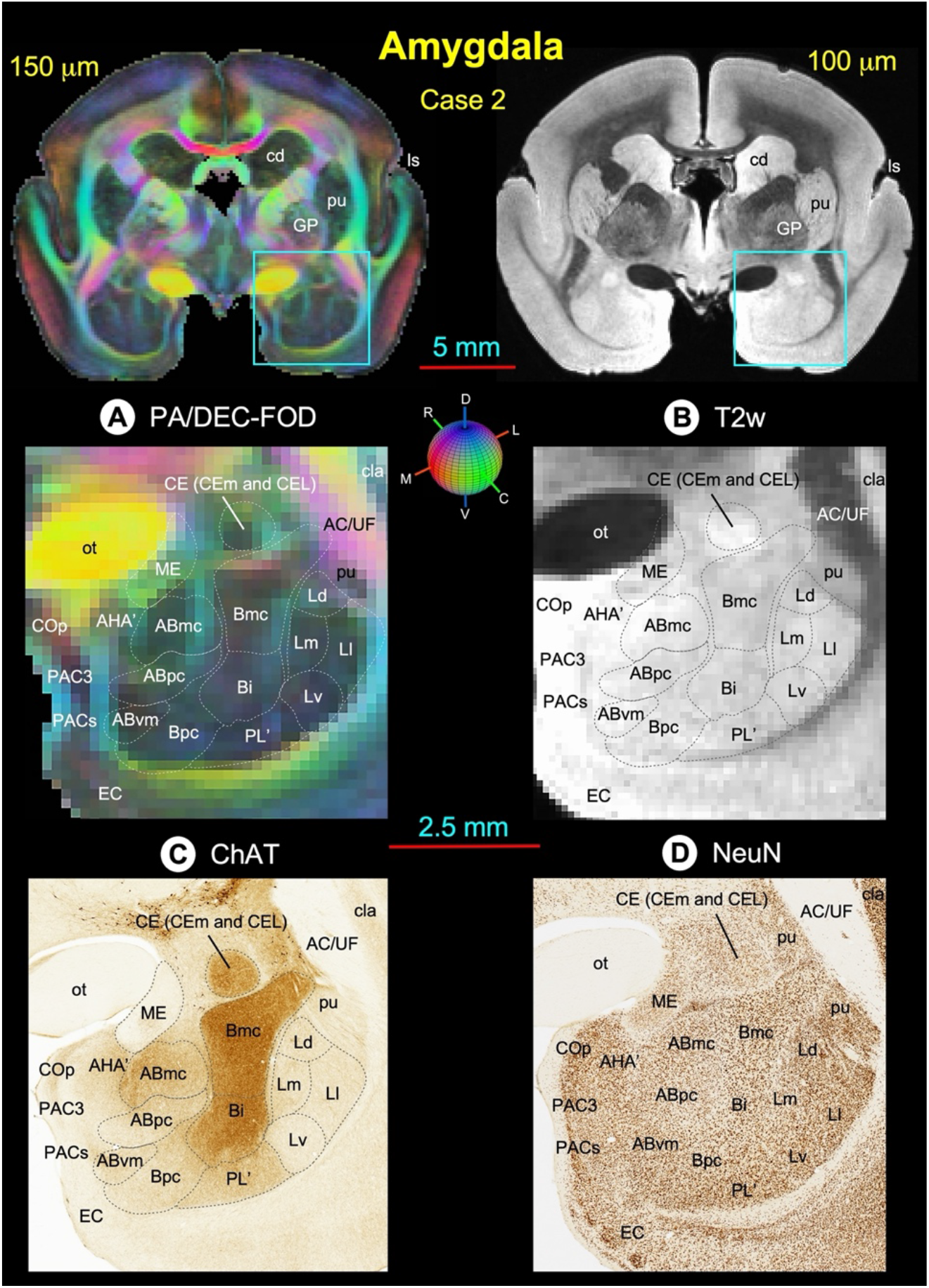
Amygdala. Zoomed-in view of the green boxed region on the top shows the spatial location of different amygdaloid nuclei and their subregions in coronal MAP-MRI (PA/DEC-FOD) and T2W images with matched ChAT and NeuN stained sections (A-D). This coronal slice corresponds to the midportion of the right amygdala. The dashed outlines of different subregions on the PA/DEC-FOD and T2W images are derived from the ChAT-stained section (C). The lateral is on the right, and the dorsal is on the top. ***Abbreviations***: ABmc-accessory basal nucleus, magnocellular subdivision; ABpc-accessory basal nucleus, parvicellular subdivision; ABvm-accessory basal nucleus, ventromedial subdivision; AC/UF-anterior commissure/uncinate fasciculus; AHA’-amygdalohippocampal area; Bi-basal nucleus, intermediate subdivision; Bmc-basal nucleus, magnocellular subdivision; Bpc-basal nucleus, parvicellular subdivision; cd-caudate nucleus; CE-central nucleus; CEL: central nucleus, lateral division; CEm-central nucleus, medial division; Cop-posterior cortical nucleus; cla-claustrum; EC-entorhinal cortex; GP-globus pallidus; Ld-lateral nucleus, dorsal subdivision; Ll-lateral nucleus, lateral subdivision; Lm-lateral nucleus, medial subdivision; ls-lateral sulcus; Lv-lateral nucleus, ventral subdivision; ME-medial nucleus; ot-optic tract; PAC3-periamygdaloid cortex 3; PACs-periamygdaloid cortex, sulcal portion; PL’-paralaminar nucleus; pu-putamen. ***Orientations:*** D-dorsal; V-ventral; R-rostral; C-caudal; M-medial; L-lateral. Scale bar: 5 mm (top images); 2.5 mm (A-D).

### Marmoset *ex-vivo* and in-vivo comparison (Fig. 16)

**Fig. 16.**
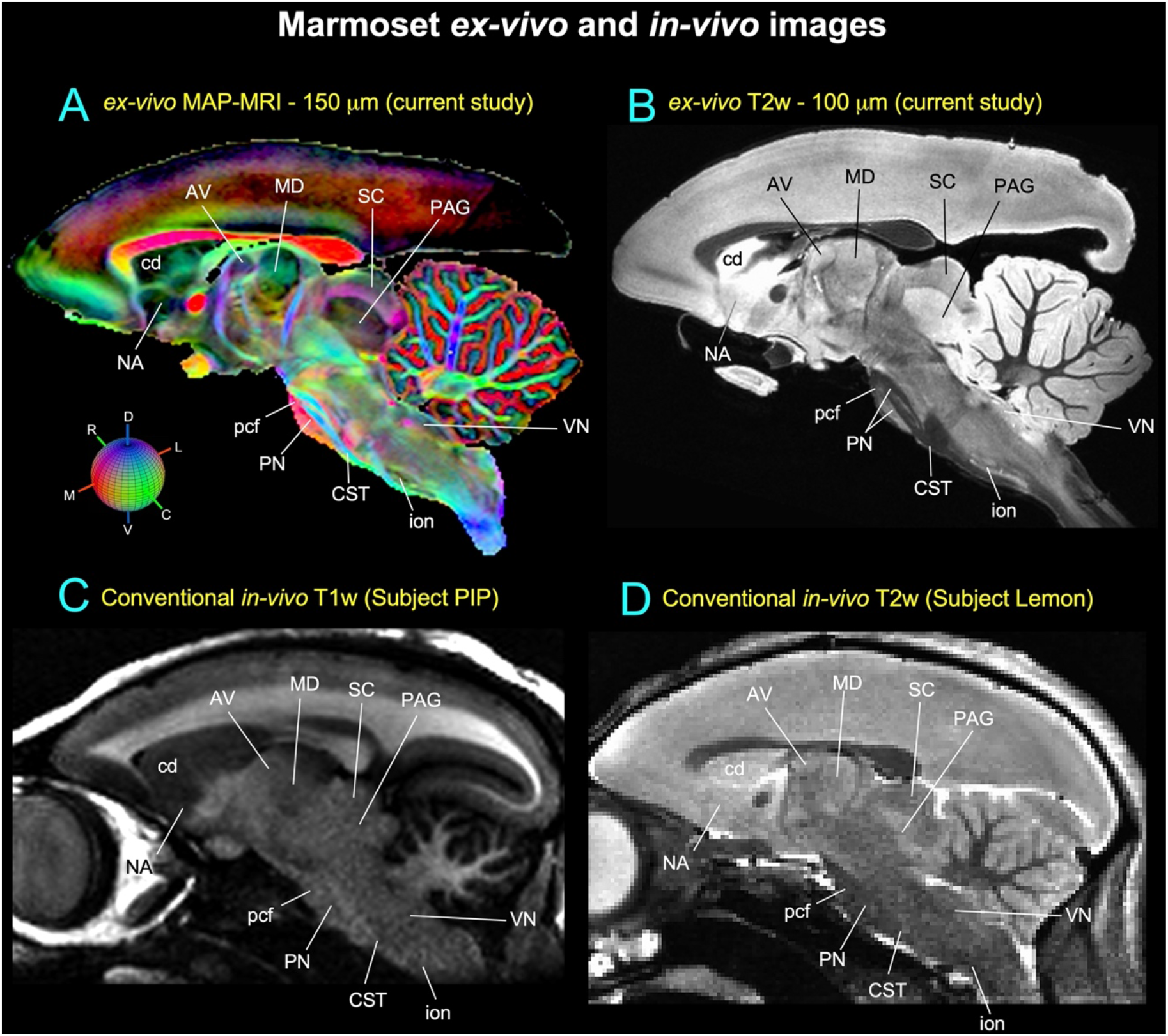
Subcortical regions in ex vivo and in vivo MRI. The closely matched sagittal MR slices from high-resolution *ex vivo* MAP-MRI (**A**, current study), T2W (**B,** current study), and conventional in vivo T1W and T2W (**C, D**) volumes show the selected brainstem nuclei and fiber tracts. Note that the nuclei (dark gray regions) are sharply delineated from the surrounding fiber bundles of different orientations, identified on the MAP-MRI with directional information (*fODF*s/DEC) and T2W images (**A, B**). In contrast, the delineations of nuclei from the surrounding white matter pathways are barely visible in conventional MRIs (**C, D**). ***Abbreviations***: AV-anterior ventral nucleus of the thalamus; cd-caudate nucleus; CST-corticospinal tract; ion-inferior olivary nucleus; MD-mediodorsal nucleus; NA-nucleus accumbens; PAG-periaqueductal gray; pcf-pontocerebellar fibers; PN-pontine nuclei; SC-superior colliculus; VN-vestibular nuclei. ***Orientations:*** D-dorsal; V-ventral; R-rostral; C-caudal; M-medial; L-lateral.

## Discussion

Despite its importance as a model for human brain development and neurological disorders, the marmoset lacks comprehensive MRI-histology-based parcellation of subcortical regions. In this study, we have described the spatial location, boundaries, and microarchitectural features of subcortical areas obtained from *ex-vivo* marmoset brains using high-resolution MAP-MRI, T2W, and MTR images at 7T (i.e., 100-150 μm resolution), aided by histology with seven different stains obtained from the same brain specimens. Our results demonstrate that, at a high spatial resolution, the MAP-MRI with fiber orientation distribution functions (fODFs) can distinguish many gray matter structures, such as sensory and motor nerve nuclei and fiber pathways of different orientations in deep brain structures. Additionally, the integrated multimodal approach using different MRI parameters and multiple histological stains enabled detailed noninvasive anatomical mapping of nuclei and their subregions in the basal ganglia, thalamus, brainstem, limbic structures, and associated fiber bundles. In contrast, the anatomical details evident in MAP-MRI and other MRI parameters are invisible in conventional T1W or T2W images (see Fig. 16). We believe our current multi-modal approach has produced a more objective and reproducible description of subcortical targets in this primate species. In the following sections, we first discuss our results along with previous studies of these deep brain targets in primates and then highlight the potential use of MAP parameters to map the deep brain structures and construct 3D digital template atlases in humans.

### Anatomical mapping of subcortical nuclei and fiber tracts at high-field MRI in primates

Our high-resolution *ex vivo* MAP-MRI, T2W, and MTR images revealed sharp borders and high contrast between gray and white matter in the deep brain structures (e.g., basal ganglia, thalamus), resulting in the precise demarcation of fine anatomical structures such as nuclei and sub-laminae, consistent with the histological findings from multiple stained sections prepared from the same brain specimens. Our fODFs-derived directionally encoded color (DEC-FOD) maps also revealed several nuclei and the orientation of small and large fiber tracts that link basal ganglia with the thalamus (e.g., lf-lenticular fasciculus, Fields of Forel H, H1, H2) and other long-range fiber bundles such as corticospinal/corticobulbar tract, pontocerebellar fibers, and medial lemniscus (Figs. 8, 12). The fields of Forel in human brains are the most recognized regions in deep brain stimulation of the subthalamic nucleus (STN) in Parkinson’s disease (Neudorfer et al., 2017). Our high-dimensional MAP (PA/DEC-FOD) and other MRI parameters revealed microarchitectural features within the basal ganglia subregions (e.g., striatum, Fig. 4) that can be comparable to the neurochemically defined compartments with distinct input and output connections called Patch/striosomes and matrix (Brimblecombe and Cragg, 2017; Crittenden and Graybiel, 2011; Eblen and Graybiel, 1995; Gerfen, 1989; Graybiel and Ragsdale, 1978; Hirsch et al., 1989; Martin et al., 1993; Smith et al., 2016).

Our high-dimensional MTR and MAP-MRI images showed several laminae separating subcortical gray matter regions in the basal ganglia and thalamus. The lateral and medial medullary laminae (lml and mml, respectively) are located between the putamen and pallidum (Gpe and GPi), and the internal medullary lamina (iml) is located between the anterior and lateral thalamic groups. The lml and mml in the marmoset and the macaque (Saleem et al., 2021) are comparable to the lamina pallidi lateralis and lamina pallidi medialis, respectively, in humans visualized in susceptibility-weighted imaging (SWI) at 7T *in vivo* (Abosch et al., 2010; Schaltenbrand and Wahren, 1977). The third lamina (lamina pallidi incompleta) subdivides the internal segment of the globus pallidus (GPi) into lateral and medial divisions was also found in the human SWI (Abosch et al., 2010). Although this lamina is absent or less prominent in marmoset and macaque monkeys, the lateral and medial subregions of the GPi in marmoset are distinguished based on the orientation of neuronal processes in our parvalbumin stained sections (magnified regions in Fig. 5E).

The location of basal ganglia subregions, such as striatum (caudate and putamen), pallidum (GPe and GPi), substantia nigra, and subthalamic nucleus, and associated fiber bundles as described above in marmosets is comparable to macaque monkeys and humans (Deistung et al., 2013a; Saleem et al., 2021; Straub et al., 2019; Tang et al., 2018). In humans, many of the deep brain structures were mapped *ex vivo* or *in vivo* with ultrahigh-resolution MRI using multiple anatomical contrasts (Abosch et al., 2010; Deistung et al., 2013a; Deistung et al., 2013b; Keuken et al., 2014; Lenglet et al., 2012). These subcortical structures represent common targets for deep brain stimulation (DBS) (for review, see (Plantinga et al., 2014) for treating cognitive dysfunction in patients with motor-related symptoms in Parkinson’s disease, traumatic brain injury (TBI), and neuropsychiatric disorders (Kundu et al., 2018; Sullivan et al., 2021; Wichmann and Delong, 2011). Together, MAP-MRI and other MRI contrast like T2W could facilitate accurate delineation of the potential DBS targets in the NHP model of Parkinson’s disease and other neurological disorders (Xiao et al., 2016). We also identified the medial and lateral subregions of the habenular complex based on the MR contrast in MAP-MRI (DEC-FOD) and staining differences of neuropil in the corresponding histological sections stained with PV and SMI-32 (Fig. 11G-I). The distinction between the medial and lateral parts of the habenular complex observed in the marmoset can be comparable to human and macaque habenular complex (Diaz et al., 2011; Saleem et al., 2021).

We show that high-resolution MAP-MRI and T2W scans provide new anatomical details and directional information (fODFs/DEC) that can complement multiple histological stains. Collectively, these are crucial in delineating nuclei and fiber tracts of different sizes and orientations in deep brain structures (e.g., pontocerebellar fibers from the pyramidal tract, medial lemniscus, and brainstem nuclei in midbrain, pons, and medulla; figs. 12-14). Many of these deep brain targets and their subregions are less prominent or spatially not distinguishable from neighboring structures with a conventional T1W or T2W MRI (Fig. 16). The most important unique feature of our subcortical mapping is the strict adherence to an MRI scan with the adjacent and matched histology sections with multiple stains from the same brain. As a result, the alignment accuracy between areal boundaries and gross anatomical features is optimized for identifying region-of-interest in this study (e.g., Figs. 11, 13, 14, 15).

### Marmoset versus macaque

In some subcortical regions, we found distinct MR signal intensity differences between the marmoset and macaque monkeys. For example, the subregions of the basal ganglia, such as the globus pallidus and the ventral pallidum, and the deep cerebellar nuclei exhibited significantly more hypointense signals in the T2W images of macaque than in the marmoset. The MRI studies in humans and nonhuman primates have confirmed and interpreted this hypointense signal to a high level of intracellular iron deposit (Bizzi et al., 1990; Drayer et al., 1986; Hardy et al., 2005; Manova et al., 2009). We confirmed this finding in the matched histology sections derived from the same macaque and marmoset brain specimens stained with Perls Prussian blue stain (Figs. 9, 10). In the macaque, the neuropil of the basal ganglia subregions and the deep cerebellar nuclei stained intensely with Prussian blue confirms the presence of a high-level iron. The spatial location of these intensely stained regions matched well with the hypointense signal intensity found in these deep brain structures in macaque T2W images. In contrast, these subregions in the marmoset revealed very faint staining with Prussian blue (Fig. 10). Based on the weak neuropil staining with Prussian blue and significantly less hypointense signals in T2W images, we conclude that these deep brain structures in the marmoset contain less iron than in the macaque monkey. We also found other differences between these two species in the subcortical white matter. For example, the anterior limb of the internal capsule, cerebral peduncle, anterior commissure, and the deep cerebellar white matter exhibited more hypointense signal in the marmoset than in the macaque (Fig. 9).

Our histochemical and immuno-histochemical stained sections in the marmoset also revealed rostral and caudal subregions of the red nucleus with different chemoarchitectonic features (Fig. 7). These subregions show discrete borders and more hypointense signal in the marmoset T2W images than in the macaque (Saleem et al., 2021) (their Fig. 7). Similar to marmoset, the red nucleus demonstrated a clear boundary with a significantly decreased (hypointense) signal in T2- and susceptibility-weighted MRI in healthy human subjects (Abosch et al., 2010) and T2W images in postmortem human brain specimens (Massey et al., 2012). Thus, the T2W MRI marker is unsuitable for delineating the red nucleus in macaque monkeys.

### Future direction: MAP-MRI-based mapping and atlases of deep brain structures in humans

MAP-MRI has been applied to numerous studies using healthy volunteers (Avram et al., 2016), aging subjects (Bouhrara et al., 2023), and patient populations with Parkinson’s Disease (Le et al., 2020), multiple sclerosis (Boscolo Galazzo et al.), epilepsy (Ma et al., 2020), ischemic stroke (Boscolo Galazzo et al., 2018), brain injury and cancer (Jiang et al., 2021). In this study, we mapped subcortical structures in the marmoset monkey brain using MAP-MRI data acquired with very high-resolution *ex vivo*. We further illustrated how our multimodal approach could locate and segment small subcortical structures using MRI and histology. These results suggest the utility of high-resolution mapping in studies of marmoset monkey disease models and highlight the potential of high-resolution MAP-MRI in delineating small subcortical structures based on differences in microstructural properties. The spatial resolution of MAP-MRI data acquired on clinical scanners is significantly lower, approximately 1-2 mm. Nevertheless, promising advances in the RF coil engineering (Keil et al., 2013; Truong et al., 2014), gradient coil design (McNab et al., 2013), dMRI pulse sequence design (Avram et al., 2014b) and spatial encoding (Feinberg et al., 2010; Setsompop et al., 2018) is expected to significantly improve the spatial resolution and signal-to-noise ratio to allow submillimeter clinical MAP-MRI scans in the near future (Huang et al., 2021). Concurrently, new clinically feasible diffusion encoding strategies (Avram et al., 2010; Avram et al., 2019; Avram et al., 2021) and analyses (Avram et al., 2022a; Magdoom et al., 2023) are being developed to quantify specific microscopic tissue water pools without the need to increase the spatial resolutions. Taken together, these advances will enable the construction of high-resolution cortical and subcortical maps and atlas templates of the human brain that will improve localization of fMRI responses, neurosurgical navigation, high-precision placement of recording and stimulating electrodes in patients with Parkinson, mild TBI (mTBI), epilepsy, and other diseases.

### Summary and Conclusion

Using combined high-resolution MAP- and other MRI parameters and corresponding histological sections with multiple stains, we have mapped the spatial locations, boundaries, and micro-architectural features of subcortical gray and white matter regions in marmoset monkeys *ex vivo*. The strengths of this work compared to others are that 1) it uses an extensive range of high-resolution MRI parameters with fODFs/DEC acquired at 7T, 2) it utilizes both high-resolution MRI with 100-150 μm resolution and corresponding histology information from the same brain specimen to map anatomical structures in depth, and 3) provides a complete, accurate, and detailed mapping of subcortical structures, including the brainstem with the cerebellum. Moreover, the methodological consistency with our previous study in the macaque brain (Saleem et al., 2021) allows us to draw meaningful conclusions between species-related differences in the structure and organization of deep brain nuclei. The high-resolution mapping of subcortical regions using multiple MRI contrasts, combined and correlated with histology, can reveal previously invisible structures radiologically, providing a readily usable anatomical standard for region definition of subcortical targets in the marmoset. This multi-modal mapping offers a roadmap toward identifying salient brain areas *in vivo* in future neuroradiological studies. Tracing and validating these critical brain regions in 3D are imperative for neurosurgical planning, navigation of deep brain stimulation probes for DBS, and establishing brain structure-function relationships.

## Credit author statement

### Kadharbatcha S. Saleem

Corresponding author, designed the study, coordinated the project, prepared the specimens for MRI and histology; mapped, segmented, and verified all the anatomical regions of interest with reference to MRI and histology; made illustrations; wrote, edited, and streamlined the manuscript.

### Alexandru V Avram

Conducted MRI experiments, including MAP-MRI scans; processed and analyzed all MRI data; helped with atlas data registration; and wrote and edited the manuscript. ***Cecil Chern-Chyi Yen:*** Conducted MRI experiments and commented on the manuscript.

### Kulam Najmudeen Magdoom

Conducted MRI experiments and commented on the manuscript. ***Vincent Schram:*** Obtained high-resolution images of all histology-stained sections in the microscope imaging center (MIC) at NICHD and commented on the manuscript.

### Peter J Basser

helped design the study, edited the manuscript, and provided research resources.

## Acknowledgments

This work was supported by the CNRM Neuroradiology/Neuropathology Correlation/Integration Core, 309698-4.01-65310 (CNRM-89-9921); Intramural Research Program of the *Eunice Kennedy Shriver* National Institute of Child Health and Human Development; the Intramural Research Program of the National Institute of Neurological Disorders and Stroke; “Connectome 1.0: Developing the next generation human MRI scanner for bridging studies of the micro-, meso -and macro-connectome “, NIH BRAIN Initiative 1U01EB026996-01. We thank James Pickel, Transgenic core facility at NIMH, for providing perfusion-fixed marmoset brain specimens for our experiments; Michal Komlosh for preparing the brain specimen for MRI and helping with the MRI data collection. We also thank Cirong Liu for providing us with the standard control in vivo T1- and T2W MRI volumes and high-resolution diffusion MRI dataset of marmoset monkeys. Finally, we thank the Microscope imaging core (MIC) at NICHD for help with the high-resolution imaging of histology sections. Dr. Du and his team at FD NeuroTechnologies in Columbia, Maryland, for all histological processing of the brain tissue.

---

**Supplementary Figure 1.**
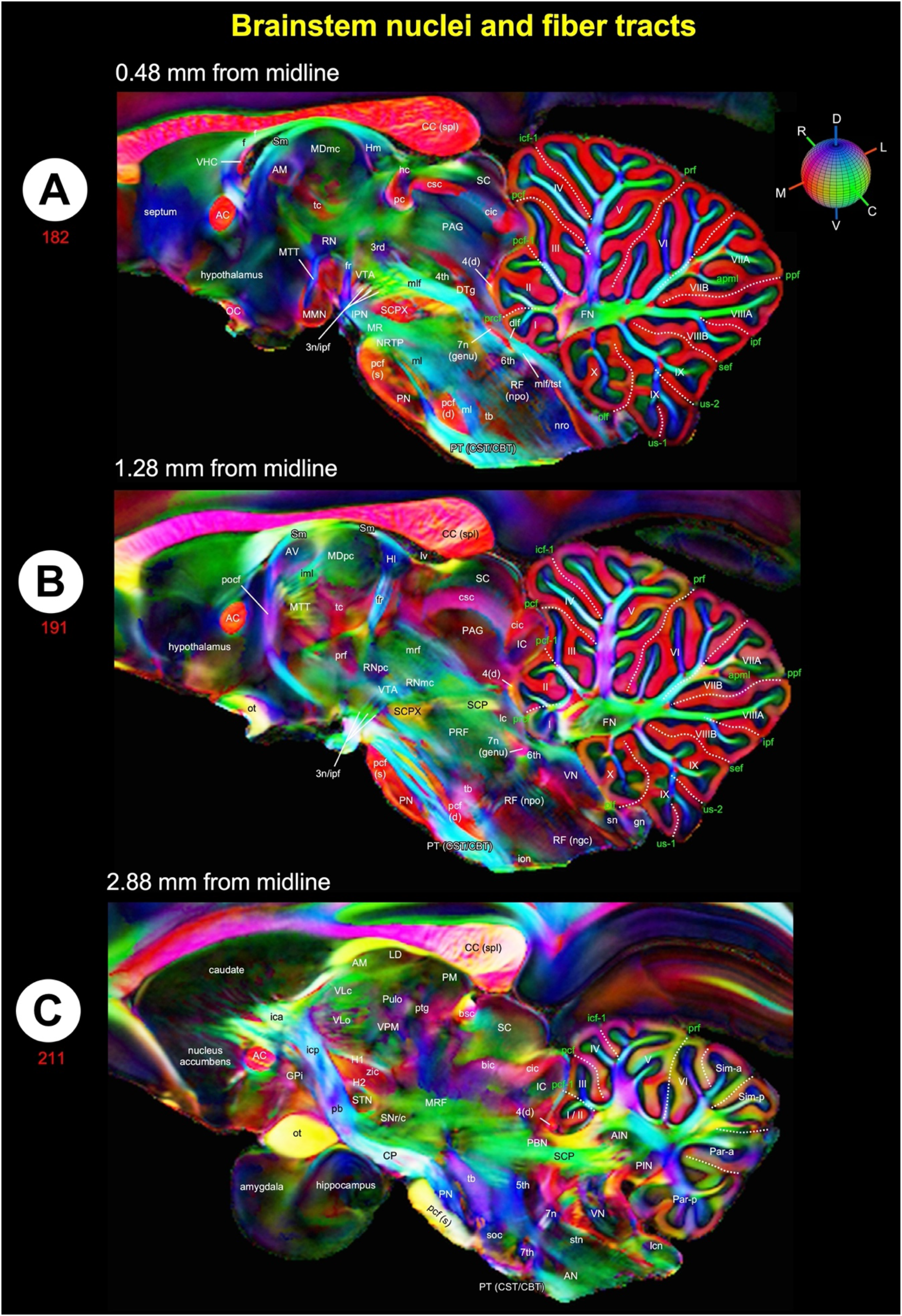
Mapping of Brainstem nuclei and fiber tracts using ultra-high-resolution dMRI (80 μm) **(A-C)** Mediolateral extent of the brainstem (midbrain, pons, and medulla) with the cerebellum in three sagittal DEC-FOD images, spaced 0.48 mm, 1.28 mm and 2.88 mm from the midline. We mapped brainstem nuclei and fiber tracts on these DEC-FOD (80 μm resolution) sagittal slices obtained from the publicly available dataset (Liu et al., 2020) and with permission from the author. This brain specimen did not include the entire medulla in the brainstem. For the complete brainstem data, including the midbrain, pons, and medulla, see Fig. 12. ***Brainstem***: white or black letters indicate the gray matter regions or nuclei, and the major fiber tracts run in different directions in the brainstem (see the color-coded sphere with directions at the top in A). Red, green, and blue in these images indicate the anisotropy along mediolateral (ML), rostrocaudal (RC), and dorsoventral (DV) directions, respectively. Other gray and white matter subregions rostral to the brainstem (e.g., thalamus and basal ganglia) are also included. ***Cerebellum***: white letters indicate different cerebellar lobules, green letters show the names of cerebellar sulci and fissures, and white dashed lines show the sulci separating different lobules. ***Abbreviations for Brainstem and surrounding regions (A-C):*** 3^rd^-oculomotor nuclei; 3n-oculomotor nerve; 4^th^-trochlear nuclei; 4(d)-trochlear nerve decussation; 5^th^-trigeminal nuclei; 6^th^-abducent nuclei; 7^th^-facial nuclei; 7n-facial nerve; 7n (genu)-genu of the facial nerve; AC or ac-anterior commissure; AM-anterior medial nucleus; AN-ambiguous nucleus; aq-aqueduct; AV-anterior ventral nucleus; bic-brachium of inferior colliculus; bsc-brachium of the superior colliculus; CBT-corticobulbar tract; CC (spl)-splenium of corpus callosum; cic-commissure of inferior colliculus; cn-cuneate nucleus; CP-cerebral peduncle; csc-commissure of superior colliculus; CST-corticospinal tract; dlf-dorsal longitudinal fasciculus; DR-dorsal raphe; dsct-dorsal spinocerebellar tract; DTg-dorsal tegmentum; dttt-dorsal trigemino thalamic tract; f-fornix; fr-fasciculus retroflexus; gn-gracile nucleus; GPi-globus pallidus internal segment; H1-H1 field of Forel; H2-H2 field of Forel; hc-habenular commissure; Hl-lateral habenular nucleus; Hm-medial habenular nucleus; IC-inferior colliculus; ica-anterior limb of the internal capsule; icp-posterior limb of the internal capsule; iml-internal medullary lamina; ion-inferior olivary nucleus; ipf-inter peduncular fossa; IPN-interpeduncular nucleus; lc-locus coeruleus; lcn-lateral cuneate nucleus; LD-lateral dorsal nucleus; lv-lateral ventricle; MDmc-medial dorsal nucleus, magnocellular division; MDpc-medial dorsal nucleus, parvicellular division; ml-medial lemniscus; mlf-medial longitudinal fasciculus; MMN-medial mammillary nucleus; MR-median raphe; mrf-mesencephalic reticular formation; MRF-midbrain reticular formation; MTT-mammillothalamic tract; NRTP-nucleus reticularis tegmenti pontis; OC-optic chiasm; ot-optic tract; PAG-periaqueductal gray; pb-pontine bundle; PBN-parabrachial nucleus; pc-posterior commissure; pcf (d)-pontocerebellar fiber, deeper part; pcf (s)-pontocerebellar fibers, superficial part; PM-medial pulvinar; PN-pontine nuclei; pocf-postcommissural fornix; PRF-pontine reticular formation; prf-prerubral field; PT-pyramidal tract; ptg-posterior thalamic group; Pulo-pulvinar oralis nucleus; RF (ngc)-reticular formation, nucleus gigantocellularis; RF (npo)-reticular formation, nucleus pontis centralis oralis; RN-red nucleus; RNmc-red nucleus, magnocellular division; RNpc-red nucleus, parvicelluar division; SC-superior colliculus; SCP-superior cerebellar peduncle; SCPX-superior cerebellar peduncle decussation; Sm-stria medullaris; sn-solitary nucleus; SNr/c-substantia nigra pars reticulata/compacta; soc-superior olivary complex; stn-spinal trigeminal nucleus; STN-subthalamic nucleus; tb-trapezoid body; tc-thalamic commissure; VHC-ventral hippocampal commissure; VLc-ventral lateral caudal nucleus; VLo-ventral lateral oral nucleus; VN-vestibular nuclei; VPM-ventral posterior medial nucleus; VTA-ventral tegmental area; zic-zona incerta. ***Abbreviations for cerebellar regions (A-C):*** I to X-cerebellar lobules; AIN-anterior interposed nucleus; FN-fastigial nucleus; Par_a-paramedian lobule anterior part; Par_p-paramedian lobule posterior part; PIN-posterior interposed nucleus; Sim_a-anterior part of the simple lobule; Sim_p-posterior part of the simple lobule. ***Cerebellar sulci/fissures (A-C):*** apml-ansoparamedian lobule; icf1-intraculminate fissure 1; ipf-intrapyramidal fissure; pcf-preculminate fissure; pcf1-preculminate fissure 1; plf-posterolateral fissure; ppf-prepyramidal fissure; prcf-precentral fissure; prf-primary fissure; sef-secunda fissure; us1-uvular sulcus 1; us2-uvular sulcus 2.

